# A decrease in transcription capacity limits growth rate upon translation inhibition

**DOI:** 10.1101/599183

**Authors:** Qing Zhang, Elisa Brambilla, Rui Li, Hualin Shi, Marco Cosentino Lagomarsino, Bianca Sclavi

**Affiliations:** LBPA, UMR 8113 CNRS, ENS Paris-Saclay, 94235 Cachan, France; CAS Key Laboratory of Theoretical Physics, Institute of Theoretical Physics, Chinese Academy of Sciences, Beijing 100190, China; School of Physical Sciences, University of Chinese Academy of Sciences, Beijing, China; LCQB, UMR 7238 CNRS, Sorbonne Université, 75005 Paris, France; Dipartimento di Fisica, Università degli Studi di Milano, 20133 Milano and INFN, Italy, and IFOM, FIRC Institute for Molecular Oncology, Milan, Italy; LCQB, UMR 7238 CNRS, Sorbonne Université, Paris, France

## Abstract

**Abstract:** In bacterial cells, inhibition of ribosomes by sublethal concentrations of antibiotics leads to a decrease in growth rate despite an increase in ribosome content. The limitation of ribosomal activity results in an increase in the level of expression from ribosomal promoters; this can deplete the pool of RNA polymerase (RNAP) that is available for the expression of non-ribosomal genes. However, the magnitude of this effect remains to be quantified. Here, we use the change in the activity of constitutive promoters with different affinities for RNAP to quantify the change in the concentration of free RNAP. The data are consistent with a significant decrease in the amount of RNAP available for transcription of both ribosomal and non ribosomal genes. Results obtained with different reporter genes reveal an mRNA length dependence on the amount of full-length translated protein, consistent with the decrease in ribosome processivity affecting more strongly the translation of longer genes. The genes coding for the β and β’ subunits of RNAP are amongst the longest genes in the *E. coli* genome, while the genes coding for ribosomal proteins are among the shortest genes. This can explain the observed decrease in transcription capacity that favors the expression of genes whose promoters have a high affinity for RNAP, such as ribosomal promoters.

**Importance:** Exposure of bacteria to sublethal concentrations of antibiotics can lead to bacterial adaptation and survival at higher doses of inhibitors, which in turn can lead to the emergence of antibiotic resistance. The presence of sublethal concentrations of antibiotics targeting translation results in an increase in the amount of ribosomes per cell and a decrease in the cells’ growth rate. In this work, we have found that inhibition of ribosome activity can result in a decrease in the amount of free RNA polymerase available for transcription, thus limiting the protein expression rate via a different pathway than what was expected. This result can be explained by our observation that long genes, such as those coding for RNA polymerase subunits, have a higher probability of premature translation termination in the presence of ribosome inhibitors, while expression of short ribosomal genes is affected less, consistent with their increased concentration.

## Introduction

Bacteria often encounter sub-lethal levels of antibiotics produced by other microorganisms in their environment. A decrease in growth rate in these conditions can allow a strain to survive long enough until the inhibitor is no longer present or, in some cases, until the bacteria becomes resistant to the antibiotic via the selection of pre-existing mutations or an increase in mutation rates (1–4). However, the mechanistic details of these response pathways often remain to be described. The cellular response to the limitation of translation activity is thought to be related to the pathway involved in the stringent response, the regulatory mechanism that decreases ribosome production in response to a decrease in amino acid availability. This is mediated by the change in concentration of the secondary messenger molecule, (p)ppGpp, that is produced by the RelA enzyme when the pool of amino acids decreases and ribosomes are not loaded with charged tRNAs (5, 6). ppGpp can directly inhibit ribosome assembly (7, 8) and the activity of RNA polymerase (RNAP) at ribosomal promoters, while increasing the activity of the promoters of genes for amino acid biosynthesis (9). The transcription of ribosomal operons can use a large fraction of the free RNA polymerase pool in the cell because of the high affinity of the ribosomal promoters for the enzyme and a high frequency of transcription initiation. Therefore, the regulation of ribosomal promoter activity can be a means by which the pool of free RNA polymerase can be repartitioned between ribosomal operon transcription and non-ribosomal mRNA synthesis. This has been referred to as the “passive control” of transcription regulation (10–14).

The ppGpp dependent feedback loop also plays a role in the regulation of ribosome content as a function of growth rate (15–17). Growth rate dependent regulation of gene expression determines the allocation of cellular resources between the production of ribosomes and that of other proteins and results in a linear increase in ribosome content with increasing growth rate (15, 18–20). In richer growth media, when the amount of amino acids is higher, ppGpp levels are lower, favoring ribosome production and a higher fraction of active ribosomes (16). In poorer media it is the inverse, accumulation of ppGpp slows down the production of new ribosomes and a smaller fraction of the ribosome pool is in an active form (19).

When ribosome activity is inhibited by sublethal concentrations of antibiotics, amino acids are used more slowly and their concentration increases, which can result in a decrease in the intracellular ppGpp pool (5). The cellular response, as predicted by the ppGpp feedback loop, is to produce a higher amount of ribosomes and an increased translation rate, however, despite this increase, the cell’s growth rate is reduced (18, 19). This has been proposed to result from a decrease in the resources available for the production of non-ribosomal proteins that become limiting for cellular metabolism (18). More recent results point to a decrease in the fraction of active ribosomes to explain the decrease in the total protein production rate (19).

To measure the effect that the inhibition of ribosome activity can have on gene expression resulting from a possible repartition of RNAP, we have compared the activity of a ribosomal promoter to that of constitutive promoters with different affinities for RNAP. This approach stems from a well-established protocol developed by Hans Bremer and coworkers of using quantitative measurements of changes in constitutive and ribosomal promoter activity as reporters of changes in the amount of free RNAP and of ppGpp (21–24).

In parallel, we analyzed transcriptomics and proteomics data from the literature on the direct and indirect effects of changing ppGpp concentration and translation limitation on gene expression (25, 26). The results from this analysis are consistent with a linear decrease with decreasing growth rate in the concentration of free RNAP available for promoter binding and transcription. We propose a model that can explain this decrease in transcriptional activity based on the observation that gene length is an important parameter on the change in protein expression in the presence of sublethal levels of chloramphenicol and that RNAP contains two of the longest gene products in the *E. coli* genome.

## Results

### Transcription regulation by ppGpp does not suffice to explain gene expression changes with increasing translation inhibition by chloramphenicol

In order to measure the effect of increasing ribosome inhibition on RNAP repartition between ribosomal and non-ribosomal promoters we have chosen three reporter cassettes. The first contains a shortened version of the well-characterized ribosomal RNA operon promoter *rrnB*P1, here called P1, that terminates at −69 from the transcription start site (27). The binding sites for Fis and the higher affinity H-NS binding site are thus omitted from this construct. This promoter has a GC-rich discriminator region at the transcription initiation site that makes the open complex sensitive to changes in negative supercoiling and to inhibition by ppGpp (Table S1) (23, 28, 29). The second promoter used here is a constitutive promoter, P5, that has consensus −10 and −35 sequences and no discriminator region. The third is PLtet, also a strong constitutive promoter with no discriminator region but with a lower affinity for RNA polymerase due to a non-consensus −10 sequence (30). Bremer and coworkers have shown that the activity of the *rrnB*P1 promoter is proportional to the concentration of ppGpp *in vivo* (23), and that the activity of constitutive promoters can be used to estimate the amount of free RNA polymerase in the cell (21, 23). Each of these promoters was placed upstream of the *gfpmut2* gene and this cassette was inserted in the chromosome together with a kanamycin resistance gene expressed divergently from the chosen promoter (Fig. S1). Growth of these strains in a 96-well plate allowed us to measure the changes in growth rate, the GFP concentration and the resulting GFP production rate (Gpr) as a function of chloramphenicol concentration (Figs. 1 and S2). We compared four different growth media, M9 with glucose, M9 with glycerol and these two media supplemented with casamino acids (cAA). This results in four different growth rates. Furthermore, it has been already shown that cells growing in a growth medium containing amino acids have a lower concentration of ppGpp (16, 23), allowing us to compare the effects due to changing concentrations of this key metabolite without the use of mutant strains that can result in secondary effects on cell metabolism due to the multiple targets of ppGpp (8, 31).

**Figure 1.**
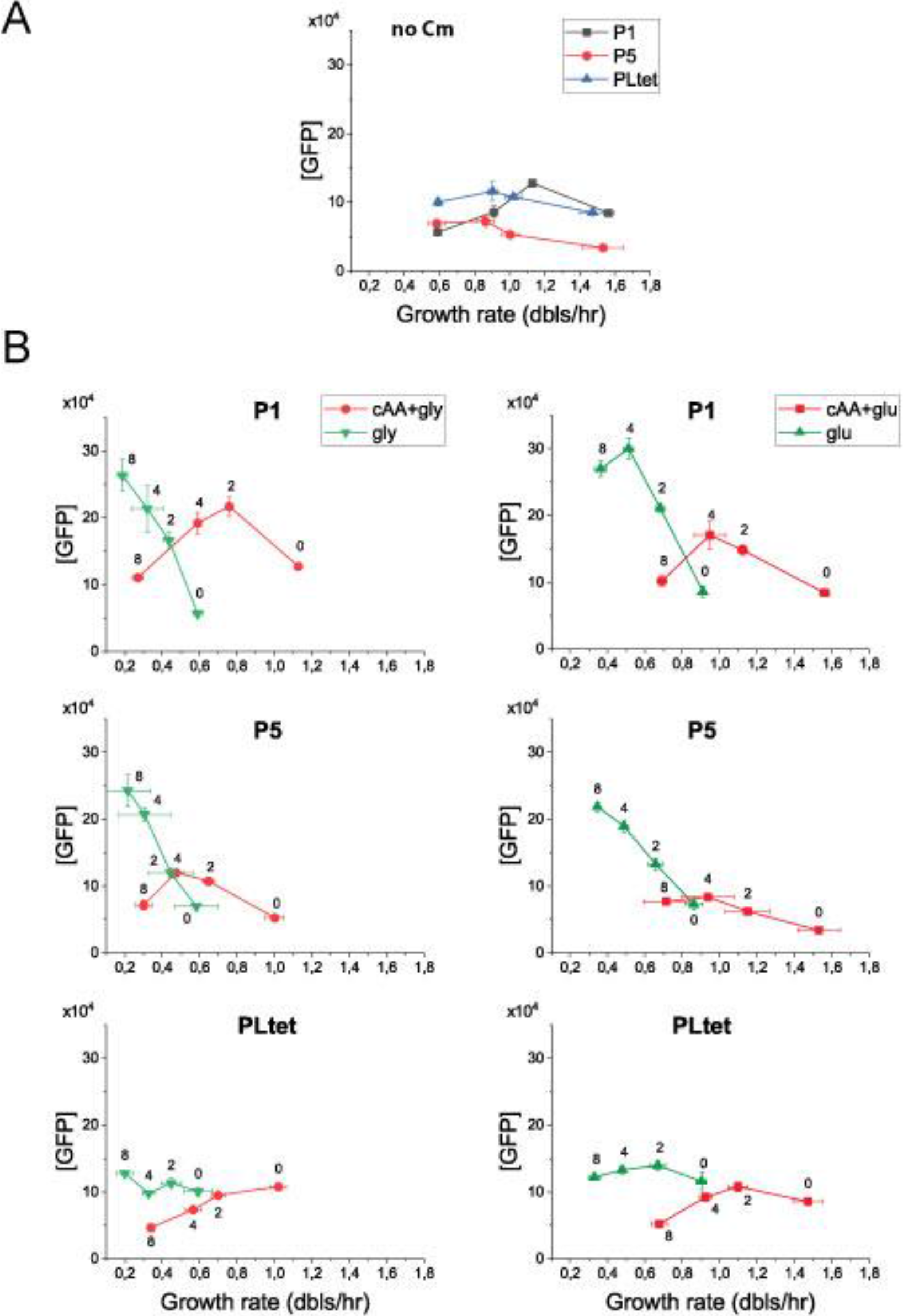
Promoters with different affinities for RNAP and different regulation by ppGpp react differently to translation limitation. **A.** Change in GFP concentration (RFU GFP/OD_600_) as a function of growth rate. Four growth media were used, from the slowest to the fastest: M9-glycerol, M9-glucose, M9-glycerol+casaminoacids, M9-glucose+casaminoacids. P5 and PLtet are both constitutive promoters with different affinities for RNAP while P1 is a shortened version of the rrnBP1 ribosomal RNA promoter with a RNAP affinity similar to P5 but regulated by ppGpp. **B.** Change in GFP concentration as a function of increasing concentration of chloramphenicol in the four growth media. The growth media with casamino acids are in red the ones without casamino acids are in green. The four points correspond to 0, 2, 4 and 8 μM final chloramphenicol concentration as noted next to the data points. The error bars represent the SEM from 3 independent experiments. The error bars smaller than the size of the symbols are not shown. The corresponding changes in GFP production rate are shown in Fig. S2. Comparison of the panels shows that ppGpp regulation at the transcriptional level alone cannot account for the change in GFP expression in response to translation limitation.

In the absence of translation inhibition, the change in promoter activity measured as a function of growth rate is consistent with previous measurements on constitutive promoters and *rrnB*P1 derived promoters (11, 27, 32) (Fig. 1A). The concentration of GFP from the constitutive promoters tends to decrease at the faster growth rates due to their lack of specific growth rate dependent regulation and the increased dilution rate (33), while the concentration of GFP expressed from the rrnBP1 promoter increases with growth rate until the last point, where Fis activation, absent in this construct, has been shown to be required for continued increased expression (27). The PLtet promoter has a lower affinity for RNAP than P5 does (see below), however when RNAP binds at the PLtet promoter it initiates transcription with a higher frequency than at P5 (30), resulting in a higher promoter activity (Fig. S2) and consequently a higher GFP concentration (Fig. 1A).

Previous work has shown that as the concentration of chloramphenicol is increased, the total RNA content relative to the total protein mass increases -reflecting the increase in ribosomal RNA- and the concentration of a reporter protein expressed from a constitutive promoter decreases (18). Therefore, the expected result here is that the GFP concentration from a ribosomal RNA promoter (P1) should increase while the concentration from a constitutive promoter (P5 or PLtet) should decrease, with the lower affinity constitutive promoter (PLtet) decreasing at a faster rate if increasing amounts of RNAP are being used for transcription of ribosomal operons. The comparison of P1 and PLtet agrees with this prediction. Unexpectedly, however, the patterns of the change in GFP production rate and GFP concentration for the P1 and P5 promoters are very similar (Figs 1 and S2). The pattern of the change of expression for these two promoter constructs depends strongly on whether the growth medium contains casamino acids, independently of the carbon source. In the absence of cAA the increase in GFP concentration as a function of Cm is significantly greater than in their presence (Fig. 1 and Fig S2). Increased expression from the P1 promoter in the growth media lacking cAA would be expected from a decrease in ppGpp as a result of increased amino acid pools due to ribosome inhibition; the P5 promoter on the other hand does not contain the GC-rich discriminator region and is not expected to show increased activity upon a decrease in ppGpp. The similar increase in GFP concentration of these two different promoters thus points to a stronger effect of ppGpp on GFP expression at the post-transcriptional level.

### Decrease in free RNAP concentration with increasing translation inhibition by chloramphenicol

Since the translation rate of GFP is shared by the three promoter constructs, it is possible to obtain an estimate of the magnitude of the promoter-specific effect of ppGpp, and of changes in free RNAP, on the transcription rate by measuring the ratios of GFP production rates. This operation “cancels out” the translation component of gene expression and isolates the transcription-specific effect (see SI text and legend to Fig. 2). Fig. 2A shows the change in the ratios of GFP production rates as a function of growth rate in the absence of chloramphenicol. The ratio of P1 to P5 rates increases rapidly between M9-glu and M9-cAA-gly, consistent with a lower level of ppGpp in the cells growing in the presence of cAA (23) increasing the probability of transcription initiation specifically from P1.

**Figure 2.**
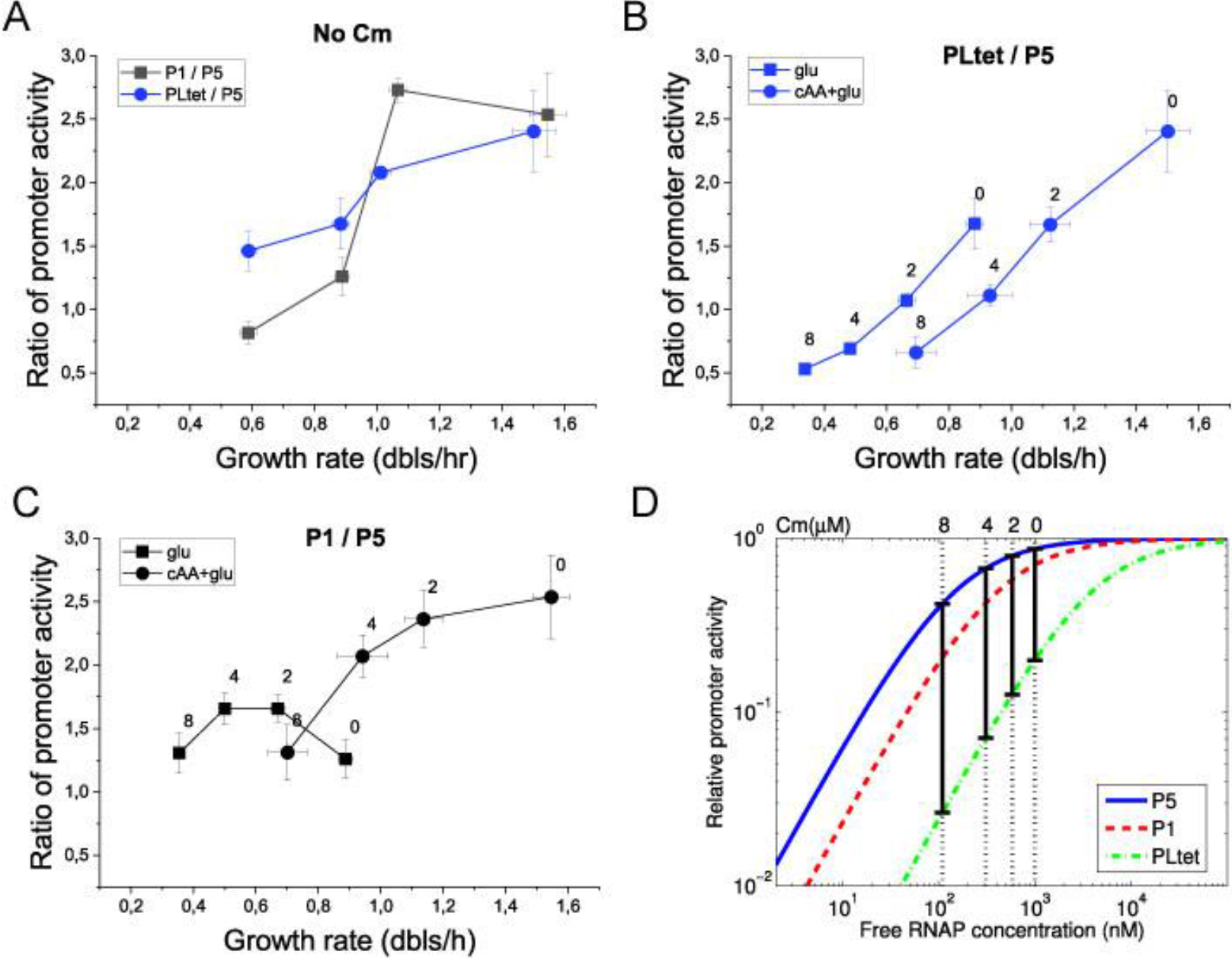
Ratios of GFP production rate for the different promoters can be used to estimate the changes in the concentration of free RNAP. **A.** Ratio of PLtet to P5 and P1 to P5 as a function of growth rate. **B.** Ratio of PLtet to P5 with increasing chloramphenicol concentration. **C.** Ratio of P1 to P5 with increasing chloramphenicol concentration. **D.** Estimated decrease in the concentration of free RNAP from the change in the ratios of promoter binding as a function of chloramphenicol concentration. To obtain this estimate, the relative activity of the promoters as a function of free RNAP concentration (c_f_) was obtained from the respective RNAP binding constants, K_i_ ∈ {K_1_, K_5_, K_Ltet_} using c_f_/(K_i_ + c_f_). The constant K_5_ is known, K_1_ and K_Ltet_ were derived by the formula K_5_exp(ΔE), where ΔE is the difference between the binding energies of RNAP with P1 (or PLtet) and P5 (see Supporting Information). The black vertical bars indicate the concentrations of RNAP that give the measured GFP production rate ratios shown in (B) for different Cm concentrations, shown above the plot. The ratio of PLtet and P5 activities for example is given by: Gpr(PLtet)/Gpr(P5) = a(K_5_ + c_f_)/(K_Ltet_ + c_f_), where a is a scaling factor. K_5_ and a were fixed by fitting the data in absence of Cm from our experiments and the literature. Data from the cells growing in cAA-glu was used so that ppGpp-dependent regulation of P1 is small and can be ignored. Fig. S3 in Supplementary Materials shows the estimation of RNAP concentration for the four growth media used. Fig.S4 shows an estimation of the change in ppGpp as a function of chloramphenicol concentration. See details of the analysis in the Supporting Information.

In the presence of Cm, the fold increase of gene expression from P1 is greater than the one of P5 in the cells that are grown without cAA, consistent with a decrease in ppGpp levels by the addition of the antibiotic (Fig. 2C). As the Cm concentration is increased further, the difference between the two promoters decreases again to the initial level. On the other hand, in the growth media with cAA, and thus lower levels of ppGpp, the P1 to P5 ratio decreases, indicating that the change in GFP production rate from P1 is lower than that of P5 as a function of increasing Cm. A similar result is also observed for the PLtet to P5 ratio, independently of the growth medium (Fig. 2B).

The comparison of two promoters with differing affinities for RNAP can be used to estimate the change in the amount of free RNAP that is available for transcription (23, 34). In the absence of Cm, the ratio of PLtet to P5 GFP production rate increases linearly with increasing growth rate (Fig. 2A), in agreement with previous estimates of the change in free RNAP as a function of doubling time (22, 33).

From the values of the RNAP affinities for these two promoters, PLtet and P5, and the relative changes in GFP production rate it is possible to estimate the change in the amount of free RNAP in the cells as a function of increasing translation limitation (see SI text). Fig. 2D and Fig. S3 illustrate this point. The RNAP binding affinity for each promoter was estimated based on a statistical-mechanical selection model developed by Berg and von Hippel (35) (see SI text). Fig. 2D shows the estimated binding curves for RNAP to each promoter; the vertical bars indicate the ratio of promoter activity corresponding to changes in the ratio measured experimentally, which were used to estimate the RNAP concentration. The values of the Cm concentration for each of these ratios is shown on the top axis. The decrease in free RNAP with increasing chloramphenicol has a stronger effect on PLtet first, then on P1 and finally on P5. This simple model of RNAP-dependent capacity can explain the data in Fig. 2B on the change in the PLtet to P5 ratio, while the P1 to P5 ratio is also influenced by changes in ppGpp concentration. Thus, using the P1 to P5 GFP production rate ratio it is possible to estimate the change in ppGpp as a function of growth rate and as a function of Cm (Fig. S4), in a similar fashion to the approach validated by Bremer and colleagues (23).

In summary, the transcriptional changes measured by the decrease in the PLtet to P5 and P1 to P5 ratios, which are in line with a decrease in free RNAP concentration, must be independent of a ppGpp-mediated repartition between ribosomal and non-ribosomal promoters, since they can also be observed in the growth media with cAA, where ppGpp levels have been previously shown to be very low (16).

### The decrease in ribosome processivity by chloramphenicol reduces the expression of longer genes more than shorter ones

In addition to the effect of RNAP availability, we hypothesized that the results observed might be dependent on the reporter gene used -*gfpmut2*, 714 bp long, or *lacZ*, 3072 bp long, coding for the β-galactosidase enzyme - since translation of longer genes would have a higher probability to end prematurely in the presence of ribosome inhibitors such as chloramphenicol acting during elongation.

In order to produce an expectation, we reasoned as follows. The processivity of translation has been shown to decrease exponentially with increasing gene length (36). If, in addition, ribosome processivity were decreased by an inhibitor, then the probability to finish the translation of a long gene would be lower compared to a shorter gene, decreasing the rate of expression of the longer gene to a greater extent. To test this hypothesis, we compared the changes in gene expression from the same constitutive promoter, P5, of two different genes, *gfpmut2*, and *lacZ*, in the presence of increasing chloramphenicol concentrations (Fig. 3A). The results show that while the concentration of GFP increases as a function of Cm concentration, β-galactosidase concentration decreases (Fig. 3A). Fig. 3B shows the change in the ratio of the shorter to the longer protein as a function of Cm concentration. The black line shows the fit obtained to a model of ribosome processivity. In the model, we derived the expression ratio of a short gene to a long gene based on the scenario where chloramphenicol hits a translating ribosome, causing the stalling of this and the following ribosomes and leading to the nonsymmetric degradation of mRNA (as depicted by Dai et al. (19), see details in SI text). It is possible to estimate the probability that a ribosome will be inhibited by the antibiotic before it reaches the end of the mRNA, P_hit_ (19). The equation can be extended to describe the dependence of P_hit_ on protein length, i.e.

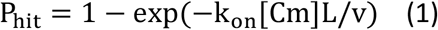

where k_on_ denotes the binding constant of Cm with ribosome (k_on_ = 0.034(μM · min)^−1^ (37)), L indicates protein length, and v is the translation elongation rate dependent on the RNA/protein mass ratio (see SI text). Using equation 1, at 8 μM Cm (the highest concentration used here) P_hit_ is 23% for LacZ (1024 aa), while for GFP it is 6% (238 aa). These results therefore indicate that a gene’s length, in addition to its promoter’s affinity for RNAP, can influence how its expression levels change in the presence of sub-lethal concentrations of ribosome inhibitors. A possible cause for the decrease in the pool of free RNAP independently of the changes in ppGpp is a decrease in the total amount of RNAP per cell. The RNA polymerase holoenzyme is composed of 5 subunits: β, β’, ω, 2 subunits of α, and a σ factor. β, β’ are among the longest genes in *E. coli* with a length of 4029 bp and 4224 bp respectively (the average gene length in *E. coli* is about 900 bp) and they could be subject to the length effect we found in our reporters.

**Figure 3.**
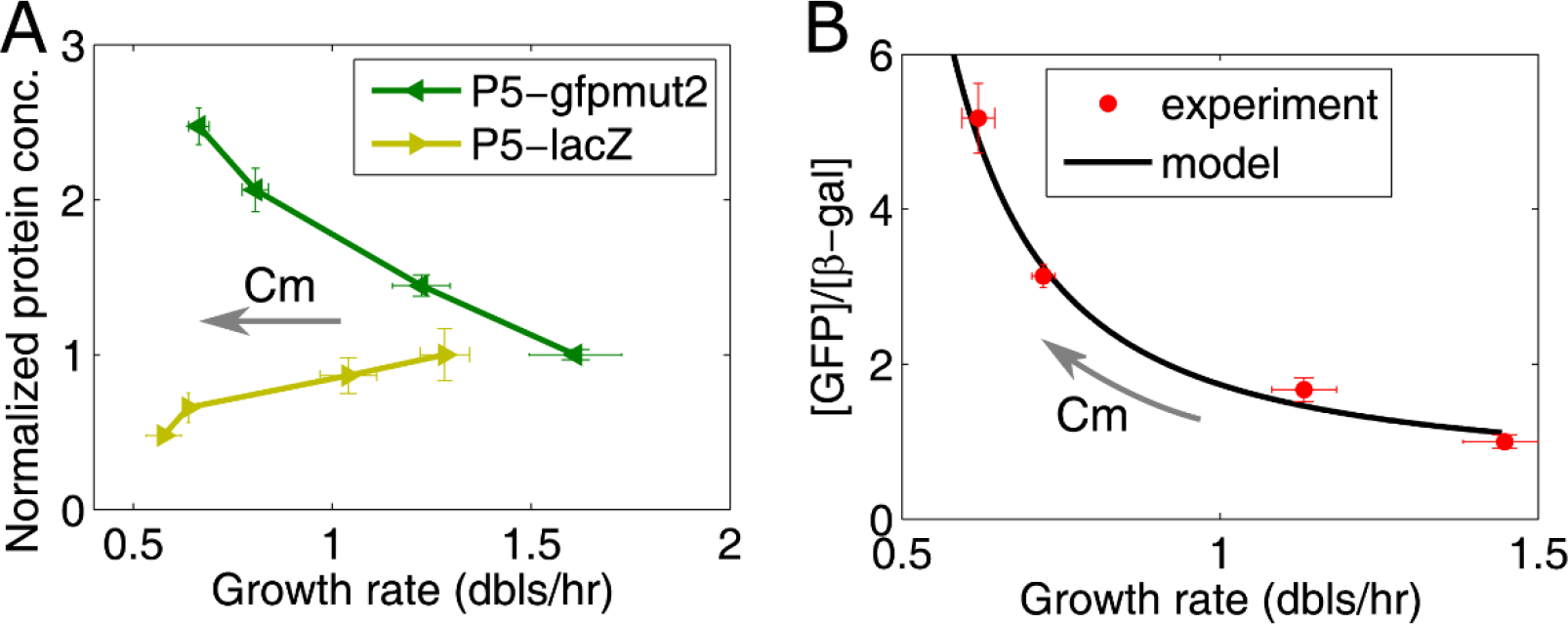
Gene length can influence gene expression under translation limitation. **A.** Change in GFP (238 aa) and β-galactosidase (1024 aa) expressed from the P5 promoter as a function of increasing chloramphenicol concentrations. The bacteria were grown in M9 glu+cAA with chloramphenicol (Cm) at concentrations of 0, 2 μM, 4 μM, or 6 μM in flasks. The concentrations were normalized by dividing the data by the point without Cm. **B.** Fit of the model to the ratio of GFP to β-galactosidase concentrations from the data in (A). The growth rate at each chloramphenicol concentration is the average over the two strains. The error bars correspond to the SEM from three independent experiments.

### A decrease in translation processivity can result in decreased expression of late operon genes

Transcription and translation are coupled via a physical interaction between RNA polymerase and the first ribosome translating the mRNA, which can be mediated by NusG or RfaH (38–41). Therefore, an additional factor that could decrease the expression of longer genes, and of late genes within an operon, in the presence of translation inhibitors is the loss of this RNAP-ribosome interaction exposing the mRNA for degradation and/or decreasing RNAP speed and processivity (42).

Previous work has shown that inhibiting ribosome activity with a higher concentration of antibiotics than was used here can result in decoupling of transcription and translation (42). To test whether these sublethal concentrations of Cm could have a similar effect on the RNAP-ribosome interaction, constructs were made where two genes of equal length coding for a red and a green fluorescent protein are placed one after the other within the same operon (Fig. 4A). In this case, there is an equal probability that a ribosome will stall during translation of either gene. However, if the first ribosome translating the upstream gene is inhibited, the one in contact with RNAP, it will also decrease the probability that the downstream gene will be transcribed by affecting the stability of the mRNA and of the transcription complex. Indeed, in these constructs we observe a decrease in the GFP to RFP ratio with increasing Cm. This could be either due to a decrease in transcription processivity from the loss of the RNAP-ribosome interaction or to an increase in the probability that the operon mRNA is degraded before RNAP finishes GFP transcription due to early termination of RFP translation, or a combination of both.

**Figure 4.**
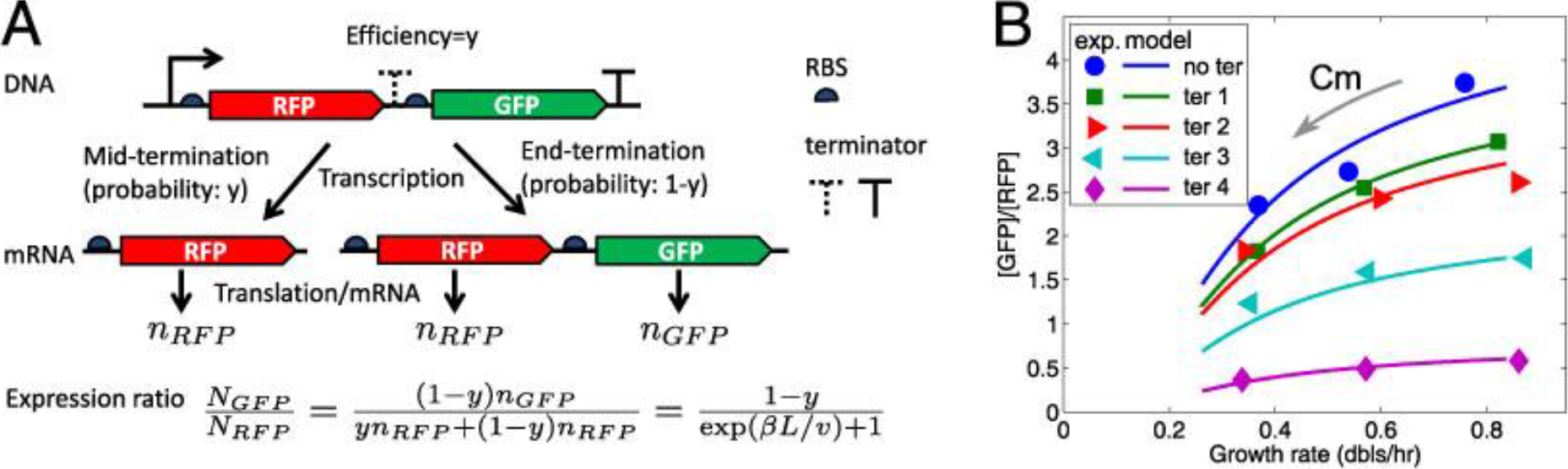
Operon position can influence gene expression under translation limitation. (**A**) Sketch for GFP and RFP fused in an operon with an intergenic terminator of efficiency y (see the details of the construction in (43)) and the formula of the expression ratio of GFP to RFP. When the efficiency y is zero, it corresponds to the case without intergenic terminator (see no ter in (B)). (**B**) The expression ratio between downstream GFP and upstream RFP (in the units of RFU(GFP)/RFU(RFP)), as a function of growth rate decreases with increasing Cm concentration. The experimental ratios (symbols) are fit with the model (lines) on Rb-stalling-induced mRNA degradation (See Supplementary Materials). ‘No ter’ and ‘ter 1-4’ correspond to terminator sequences of increasing efficiency (‘I21’ (no terminator), ‘R9’, ‘R17’, ‘W13’ and ‘R32’ respectively in (43)). The cells were grown in M9 minimal medium containing glucose. Cm was added to a final concentration of 0, 2, or 4 μM.

The formation of a terminator hairpin in the transcribed mRNA can be used to detect the presence of a ribosome-RNAP interaction (43). If a ribosome is bound on the mRNA as it is being extruded from RNAP, the terminator hairpin structure cannot fold. If inhibition of ribosome activity decouples translation from the ongoing transcription, then the RNA will be allowed to fold and transcription will stop. Transcription terminators of different strength have been inserted between the two genes (43). The efficiency of these terminators is determined by the distance between the stop codon of the upstream gene and the hairpin loop forming sequence. Termination will be less efficient when this distance is shorter. If transcription and translation are decoupled, the RNAP to ribosome distance will increase and we expect that the efficiency of the weaker terminators will increase with increasing Cm. However, we observe the same fractional decrease in GFP/RFP in all the constructs, independently of the distance from the stop codon, or of the presence of the terminator (Fig. 4 and S8). Therefore, it seems that at these low Cm concentrations, the probability of decoupling of transcription and translation is not significant enough to allow for the folding of the hairpin loop. The stalling of a ribosome however, independently of its interaction with RNAP, can result in an increased probability of mRNA degradation. In this case, the stalling of non-leading ribosomes, which are in greater number than the one interacting with RNAP and thus a more probable target, can decrease the lifetime of the operon’s mRNA, decreasing the probability that translation will be completed at the *gfp* gene. These results can be reproduced by a model using the parameters values for the probability of translation termination obtained from the comparison in GFP vs LacZ translation (Fig. 3) and the terminator strengths measured in the previous study (43) (Fig. 4B) (see SI for the details of the model).

Finally, in light of these results we have analyzed data from previously published transcriptomics and proteomics studies as a function of the presence of ppGpp and at increasing sublethal chloramphenicol concentration respectively (25, 31) and we have found that the effect of promoter affinity, gene length and operon position observed here on this set of promoters and reporter proteins can help explain the global changes in gene expression in the presence of sublethal levels of chloramphenicol (SI text and SI Figs S10 - S14).

## Discussion

### A linear decrease in transcription capacity with increased translation limitation

While the amount of free RNAP decreases with decreasing growth rate as a function of nutrient content (Fig. 2A), the decrease in the presence of increasing translation limitation, within the same growth medium, has a steeper, linear, slope (Fig. 2B and Fig. S3). The evidence provided here points to a possible cellular adaptation mechanism leading to a reduction in transcription capacity when ribosome activity is compromised. This adaptation decreases the cost of transcription of untranslated mRNAs (44) and allows for more resources to be available for the synthesis of increased amounts of ribosomes to respond to the presence of translation inhibitors. Importantly, we can quantify the contribution of the change in free RNAP to changes in transcription capacity: the comparison of two different constitutive promoters with differing RNAP affinity, PLtet and P5, shows a striking difference in the change in GFP production rate and the resulting GFP concentration with increasing ribosome inhibition (Fig. 1). The decrease in the amount of free RNAP estimated by measuring the ratio of GFP production rates from these two promoters (PLtet/P5) is about 10-fold, independently of the presence of amino acids in the growth medium (Fig. 2 and Fig. S3) and therefore of the change in ppGpp concentration (see below).

We found that, as the sublethal concentration of ribosome inhibitor is increased, the growth rate decreases linearly with the decrease in the concentration of free RNA polymerase, pointing to a possible growth-limiting role for this enzyme in these conditions (Fig. 2B). The decrease in free RNAP could be due to different factors affecting the nonspecific interactions of the enzyme with the genome (22), however, the results obtained here from the comparison of the expression of two proteins of different lengths, β-gal and GFP (Fig. 3) suggest that a decrease in the amounts of the full length protein may have a significant contribution to this effect. The RNA polymerase core contains two of the longest proteins in *E. coli*, the β and β’ subunits (Fig. S12), increasing the probability that a ribosome will stall before reaching the end of the mRNA. This interpretation is further supported by the proteomics analysis of Hui *et al*. (25). They measured the change in protein fraction of over 1000 proteins in the presence of increasing concentrations of Cm by quantitative mass spectrometry. Their results show that the β and β’ subunits of RNAP remain a constant fraction of the proteome with increasing Cm and decreasing growth rate. Since decreasing growth rate is associated with decreased protein production rate, and in these conditions ribosomes are in excess, these results are in line with RNAP playing a limiting role in determining the total rate of protein production and the cell’s growth rate. Moreover, these results can shed light on a recent study by Dai et al (19) where it was proposed that in the presence of sublethal concentrations of Cm, despite an increase in the translation elongation rate due to a higher concentration of ternary complexes, the reduction in the total protein production rate results from a decrease in the active ribosome fraction, or the fraction of ribosomes that can reach the end of a mRNA in the presence of the inhibitor (19). Here we identify RNAP as one of the genes that is likely to be most affected by the decrease in ribosome processivity due to the length of its β and β’ subunits, while shorter genes are affected to a lesser extent.

### Is the extreme length of RNA polymerase genes a feature conserved for the coupling of translation and transcription rates?

The extreme length of the β and β’ subunits of RNAP is conserved throughout bacteria (45). In the case of *Helicobacteraceae* and *Wolbachia* the two genes are even fused together (45, 46). The gene length of β and β’ subunits can vary in different strains since they are composed of independent structural modules separated by spacers of differing length (45). In *E. coli* the spacer sequences, that account for more than 25% of the total sequence, can be deleted without causing a significant decrease in transcription activity. In archea and chloroplasts some of the conserved protein modules are found in separate genes and in *E. coli* they can be split from each other to produce an active enzyme (47), suggesting that the length of these genes is not imposed by functional constraints. The reason why the RNAP and ribosomal proteins find themselves at opposite ends of the spectrum of gene lengths in bacteria is likely linked to the assembly process, structural flexibility and stability of the final multi-protein complex (48, 49); however, these results suggest that it could also play an important role for the cell’s survival in the presence of antibiotics that decrease translation processivity.

### The passive control model

Another factor that can decrease the pool of RNAP available for transcription of non-ribosomal genes is the increase in the transcription from ribosomal promoters resulting from the decrease in ppGpp, the inverse of what takes place during the stringent response and what has been referred to as the “passive control” of transcription regulation (10–14). However, the results presented here show a similar slope of the decrease in the concentration of free RNAP (Fig. 2 and S3), independently of the initial amounts of ppGpp found in the cells growing in the presence or absence of casamino acids (23). Therefore, while changes in ppGpp do affect the activity of ribosomal promoters, the magnitude of this effect is not strong enough to further decrease the activity of non-ribosomal promoters such as PLtet and P5. The main factor that partitions the limited pool of RNAP between ribosomal and non-ribosomal promoters is therefore the affinity of the specific RNAP-promoter complex formed for transcription initiation. These results can also help explain a previous report showing that the induction of persistence and of β-lactam tolerance by chloramphenicol is independent of the presence of the RelA enzyme, that, together with SpoT, is responsible for ppGpp synthesis (50).

### Change in total translation rate in response to translation limiting inhibitors

The similarity in the pattern of the change in GFP expression from the rrnBP1 and P5 promoters despite their different regulation by ppGpp indicates that upon translation limitation the main effect of ppGpp on gene expression may be via a change in the translation rate rather than the transcription rate (Fig. 1 and Fig. S15). The translation of short proteins such as GFP is barely affected by the presence of sublethal concentrations of Cm (8 % probability of drop off at the highest concentration used here) and can therefore be used to estimate the changes in total translation activity and to estimate how the translation capacity is affected by chloramphenicol challenges (Fig. S15). As the concentration of ribosome inhibitor is increased, the translation rate of GFP increases and mirrors the change of the effect of ppGpp on the P1 promoter (Fig. S15 and S4A respectively). Several factors can contribute to this increase in translation rate in the presence of Cm: the shorter than average length of ribosomal genes (Fig. S12), whose translation is not affected as strongly by translation inhibition compared to longer genes; the high affinity of RNAP for ribosomal promoters, insuring transcription despite a decrease in the concentration of free RNAP (Fig. S11); the decrease in ppGpp in the growth media lacking casamino acids (Fig. S4), resulting in an increase in both ribosomal promoter activity (Fig. 1) (5, 51) and in the fraction of active ribosomes due to the effect of ppGpp on ribosome assembly (7, 8, 19), in addition to the increase in translation elongation rate (19). The higher increase in translation rate in the growth media without cAA is consistent with a larger fraction of inactive ribosomes stored in the cells growing at these slower growth rates that can be quickly reactivated by a decrease in ppGpp concentration, not only for rapid adaptation to changes in local nutrient content, but also to respond to the presence of growth inhibitors (19, 52–55). On the other hand, in rich media, when ppGpp levels are low, a decrease in transcription rate when translation is compromised might play a crucial role for the cell’s survival, as the potential of the cell to increase its translation capacity to respond to the presence of the inhibitor is limited by the smaller fraction of inactive ribosomes (19, 56). Decoupling of transcription and translation can have several deleterious effects, including mRNA degradation and R-loop accumulation that can interfere with DNA replication, causing genome instability and increased mutation rates (57).

The decrease in growth rate due to limiting transcription is unexpected, as ribosome activity is usually thought to always be rate-limiting for bacterial growth, however, depending on the growth conditions, transcription has also been seen to become limiting in eukaryotic cells (68), pointing to different strategies of cellular adaptation to changing growth conditions and limitations. Understanding how bacteria modulate their growth rate and resource allocation in response to inhibition of growth has paramount importance in biotechnological and health applications (58–61). The results presented here provide a new cellular mechanism by which bacterial cells can decrease their growth rate in response to antibiotic stress (1, 56, 62). In summary, in the presence of sublethal concentrations of chloramphenicol, it is not translation that becomes limiting for the cell’s growth rate, or the ppGpp dependent repartition of RNAP between ribosomal and non-ribosomal promoters, but it is the decrease in total transcription capacity (Fig. 5). It remains to be established whether this is a common response to other translation limiting factors, although a similar pattern of a decrease in growth rate despite a proportional increase in ribosomal RNA content and translation rate has been observed in the past with antibiotics such as tetracycline, erythromycin, and neomycin, and limiting expression of initiator factors 2 and 3 (18, 19).

**Figure 5.**
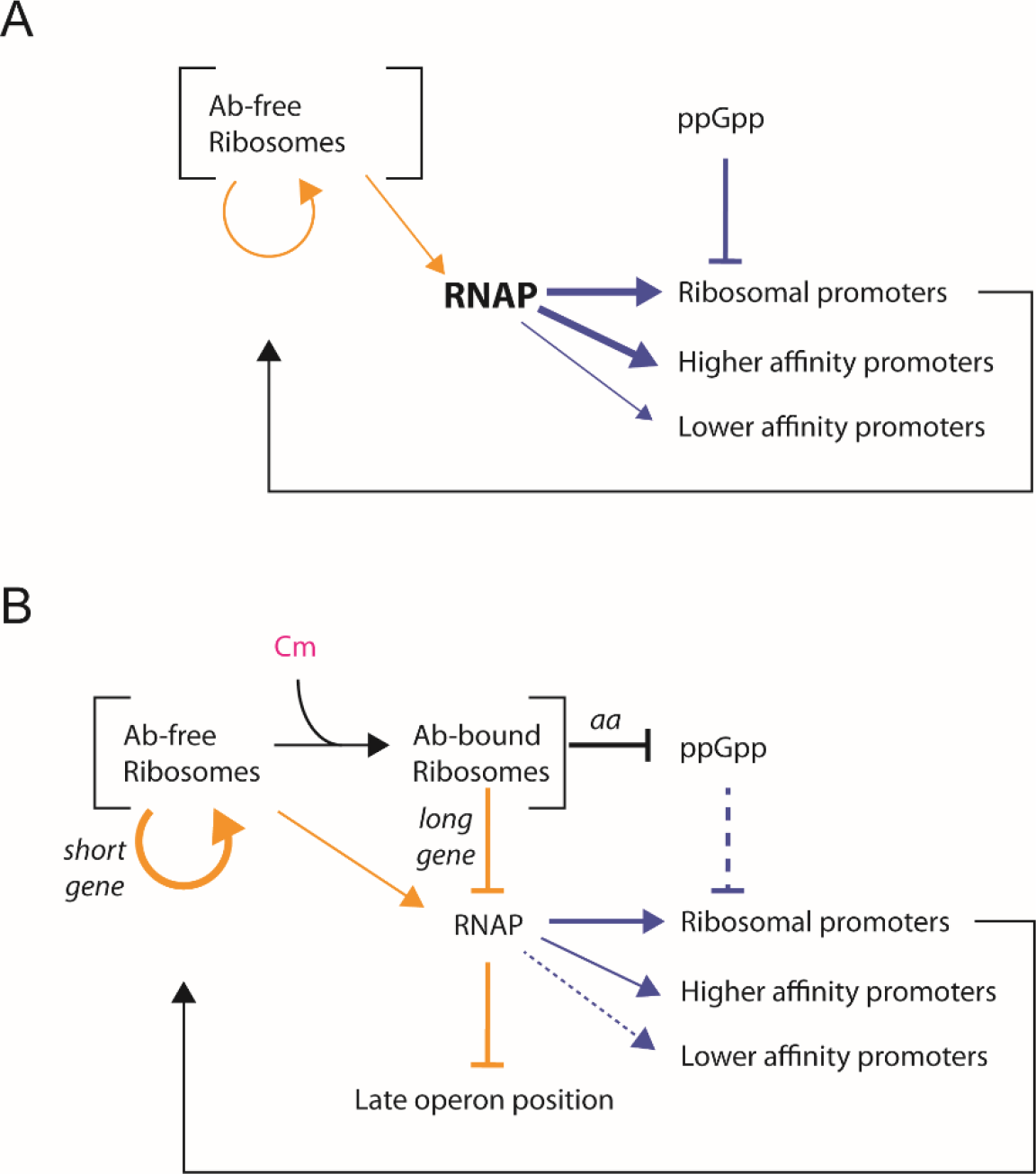
Summary. Orange arrows show translation effects and blue arrows show transcription effects. **A.** In the absence of translation limitation ribosomes translate both ribosomal and non-ribosomal genes with similar rates. The amount of RNAP available is regulated in part by the changes in ppGpp and the ensuing transcription rate of ribosomal promoters. **B.** The decrease in ribosome processivity by inhibitors such as chloramphenicol (Cm) results in translation of longer genes being prematurely terminated more frequently than translation of shorter genes. RNAP subunits β and β’ are among the longest genes in *E. coli*, while ribosomal proteins are among the shortest. The decrease in free RNAP can be measured by a decreasing ratio of high affinity to low affinity promoter genes expression rate. The decrease in ribosome processivity increases the probability of mRNA degradation, thus penalizing the expression of genes at the end of operons. In nutrient poor media, inhibition of ribosome activity by chloramphenicol increases the pool of amino acids and decreases the levels of ppGpp, increasing both ribosome production and ribosome activity. Ab: antibiotic.

## Materials and Methods

### Strains, promoters and reporters

GFP and β-galactosidase were used as the reporter proteins to measure the rate of gene expression from a specific promoter. The GFP gene used is *gfpmut2* coding for a fast-folding GFP (63). The β-galactosidase gene is a 5’-end-modified *lacZ* from the pCMVbeta plasmid (64). Comparison with the wild type *lacZ* gene shows that the additional 23 amino acids do not change the results obtained with this version of the reporter gene (data not shown). The promoters include two constitutive promoters (P5, obtained from T5 phage and PLtet, i.e. P_LtetO-1_ (30) and a shortened version of a rRNA promoter (*rrnB*P1 without the upstream Fis and H-NS sites) (Table S1). The constructs of P1-*gfpmut2*, P5-*gfpmut2*, PLtet-*gfpmut2* and P5-*lacZ* with a divergent kanamycin resistance gene were inserted in the chromosome of the BW25113 *Escherichia coli* strain. P1-*gfpmut2* and P5-*gfpmut2* (for Fig. 1) were inserted at position 258235 between the convergent *crl* and *phoE* genes, PLtet-*gfpmut2* was at position 356850 between *cynR* and *codA*, and P5-*lacZ* and P5-*gfpmut2* (for Fig. 3) were at position 1395689 between *uspE* and *ynaJ*. Genome position did not have an effect on the change in reporter gene expression as a function of Cm. The double fluorescent protein system (RFP-GFP constructs) has been described previously (43). The ribosome binding sites (RBS) i.e. Shine-Dalgarno sequences, used in above constructs are all similar to the consensus UAAGGAGGU (65). The RBS for GFP (*gfpmut2*) and β-gal (*lacZ*) is GAAGGAGAU, for RFP (*mCherry*) it is AGAGGAGAA.

### Bacterial growth and fluorescence measurements

Bacterial growth was carried out in M9 minimal growth medium supplemented with 0.5% glycerol (gly), 0.5% glucose (glu), 0.5% glycerol+0.2% casamino acids (cAA+gly) and 0.5% glucose+0.2% casamino acids (cAA+glu). The pre-culture was obtained from the inoculation of one bacterial colony in LB growth medium. After overnight growth, the seed culture was washed once with PBS and diluted 200 times with the corresponding growth medium containing a specific concentration of chloramphenicol (Cm). This culture was diluted again 200 times once it reached exponential phase. The cultures were grown in flasks, shaking at 37°C and 170 rpm, and optical density and fluorescence were measured with a plate reader (Tecan, Infinite 200Pro) every 30 - 50 min. Alternatively, the cultures were grown in a 96 well plate, with 150 μL of bacterial culture per well covered by 70μL mineral oil. The culture plate was kept at 37°C in the plate reader, shaking and measuring fluorescence and OD_600_ every 5 minutes. The auto-fluorescence measured from the wild type strain Bw25113 was subtracted from the fluorescence of the fluorescent strains at the same OD (dependent on the medium). The experimental procedure of the β-galactosidase assay followed the protocol of Zhang et al.(66) except that the bacterial strains were cultivated in flasks instead of 48-well plates(18, 67, 68). The measurement of RFP-GFP constructs followed the protocol described previously (43).

### Analysis of GFP reporter expression data

Experimental data obtained from the plate reader were analyzed with Matlab to obtain growth rate, protein concentration and protein expression rate. The pipeline is shown in Fig. S1. The window in the growth curve corresponding to the exponential growth phase was defined as a linear range between an upper and a lower threshold in the growth curve plot of log(OD_600_) versus time (the thresholds determined manually or from an automated method (69) gave similar results). Growth rate was derived from the slope of log(OD_600_) versus time in exponential phase (Fig. S2B and C). GFP concentration was derived as the slope of the plot of GFP versus OD_600_ in the exponential growth phase (Fig. S2B). β-galactosidase concentration in Miller Units was obtained by the following formula (Fig. S2C)

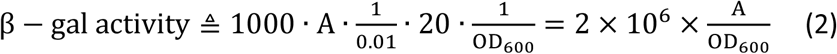

where A comes from the fit of OD_600_ as a function of time with the formula A(1 − e^−γt^)/γ and γ is a decay factor from taking into account that the reaction product o-nitrophenol is volatile(66). The rate of protein expression is defined as the product of protein concentration and growth rate (μ).

## Acknowledgments

The authors would like to thank Luca Cialdrini and Gilles Fischer for useful comments on the manuscript. This work was supported by HFSP grant RGY0070/2014 (BS, MCL and QZ) and the Strategic Pilot Project of Chinese Academy of Sciences (XDA17010504) (HS and QZ).

