## Supplementary material for "A decrease in transcription capacity limits growth rate upon translation inhibition": SI

**This PDF file includes:**

Supplementary text
Figs. S1 to S15
Table S1
References for SI reference citations

### Supporting Information Text

**Estimation of the affinity of RNAP-promoter interaction from the promoter sequence.** In 1987, Berg and von Hippel proposed a statistical-mechanical selection model to quantify the relationship between the affinity of a protein binding to a site on the DNA and individual base-pair choices in the site (1). They derived a formula to calculate the discrimination energy ($E$) for a specific sequence $\{B_l\}_{l=1}^{s}$:

$$E(\{B_l\}) = kT \sum_{l=1}^{s} \ln\left(\frac{n_{l0}+1}{n_{lB_l}+1}\right), \qquad [1]$$

where $k$ is the Boltzmann constant, $T$ is the temperature, $n_{lB_l}$ denotes the number of the sample sequences which have base $B_l$ at position $l$, $n_{l0}$ denotes the number of the sample sequences which have the cognate base at position $l$. To calculate the energy of RNAP binding to the promoter, we consider the contribution of the -10 and -35 sequences. The promoter sequences were obtained from RegulonDB (2). We consider the promoters recognized by sigma70 and the sequence of the 6 base pairs in each of the -10 and -35 regions. We obtained the frequency matrix from the literature (3), or by counting all the promoters under consideration. If we consider the bias of *E.coli* genome in base usage and the effect of the spacer length between -10 and -35 regions, equation 1 can be updated as (1)

$$E(\{B_l\}) = kT \left\{ \sum_{l=1}^{s} \ln\left[\frac{(n_{l0}+1)p_{B_l}}{(n_{lB_l}+1)p_{l0}}\right] + \ln\left[\frac{n(L_{opt})+1}{n(L)+1}\right] \right\}, \qquad [2]$$

where $p_{B_l}$ is the occurrence probability of $B_l$ in the genome, $p_{l0}$ is the occurrence probability of the cognate base at position l in the genome, $n(L)$ is the number of the promoters with the spacer length $L$ and $L_{opt}$ is the optimal spacer length. The GC content of the *E.coli* genome is 43.4%. So the occurrence probabilities of A, T, C, G in the genome are 28.3%,28.3%, 21.7% and 21.7% respectively. The dissociation constant ($K$) for one promoter (P1, P5, or PLtet) can be related to the corresponding binding energy ($E$) by $K \propto \exp(E)$.

**Estimation of the change in RNAP concentration from the analysis of the effect of Cm on GFP expression from the P5 and PLtet promoters.** P5 and PLtet are constitutive promoters, therefore their binding affinity with RNAP alone can be used to determine the probability that RNAP will be bound to the promoter at a given RNAP concentration. If the difference in RNAP affinity for two constitutive promoters is known, the difference in transcription rate from these promoters can be used to estimate the change in free RNAP concentration with Cm concentration *in vivo* (Fig. 2 in the main text and Fig. S3). In order to obtain an estimate of the absolute concentration of RNAP, the data obtained on the transcription rates in the absence of Cm can be compared to previously published values as a function of growth rate (4).

Transcription initiation can be described by Michaelis-Menten kinetics, where the process of RNAP binding with the promoter is faster than the following isomerisation steps including the formation of open complex. We formalize the relative transcription rates (TR) of P5 and PLtet as

$$TR(P5) = c_f/(c_f + K_5) \qquad [3]$$

and

$$TR(Ptet) = ac_f/(c_f + K_{tet}) \qquad [4]$$

respectively, where $c_f$ denotes the free RNAP concentration, $K_5$ and $K_{Ltet}$ are the dissociation constants for P5 and PLtet respectively, and $a$ is a scaling factor that accounts for the difference in transcription initiation frequency. The transcription initiation frequency, or promoter escape, is higher at PLtet than at P5, resulting in an increased probability of GFP expression for each binding event, and is assumed to be independent of RNAP concentration. This can explain why the GFP production rate (Gpr) PLtet to P5 ratio is greater than 1 (Fig. 2 in the main text and Fig. S3A), despite the difference in binding affinity.

The ratio of two dissociation constants ($K_j/K_i$) can be represented as an exponential function of the difference of the corresponding binding energies ($E_j - E_i$), i.e. $K_j/K_i = \exp(E_j - E_i)$. The binding energies of RNAP with the three promoters can be estimated based on their DNA sequence (see above). If one dissociation constant ($K_i$) is known, the other one, ($K_j$), can be estimated with the formula $K_j = K_i \exp(E_j - E_i)$.

We assume that the translation rate of GFP from the three promoter-*gfpmut2* constructs, including the possible effect of Cm, is the same, and that therefore it will cancel out when the ratio of GFP expression rate is taken.

The ratio of GFP production rates for PLtet and P5 is thus equivalent to the ratio of the transcription rates (TR) obtained from Eqs. 3 and 4

$$\begin{aligned} Gpr(Ptet)/Gpr(P5) &= TR(tet)/TR(P5) \\ &= a(c_f + K_5)/(c_f + K_{tet}). \end{aligned} \qquad [5]$$

If the difference in RNAP binding affinities for P5 and PLtet are known (from the free energy calculation above), the change in Gpr ratio with growth rate or with [Cm] can be used to estimate the change in free RNAP concentration (Fig. 2 in the main text).

It is also possible to estimate the absolute RNAP concentration in the cell from the available data on the change in the amount of free RNAP concentration as a function of growth rate (4) and our data on the PLtet/P5 Gpr in different growth media.

The free RNAP concentration, $c_f$, as a function of growth rate ($\mu$) can be obtained by fitting the data of Klumpp and Hwa (4), with $\log(c_f) = A\exp(-\mu_r/\mu)$, giving $A = 6.82\ \log(um^{-3})$ and $\mu_r = 0.11$ doublings/hour (Fig. S3B).

The change in $Gpr(Ptet)/Gpr(P5)$ as a function of growth rate obtained by our experimental data (Fig. S3A) is consistent with the change in RNAP concentration measured previously (4). We can thus obtain the value of $a$ and $K_5$ by fitting the Gpr ratio data with equation 5 and the RNAP concentration at the different growth rates: $a = 10.5$ and $K_5 = 90\mu m^{-3}$.

Finally, we can use equation 5 to estimate the free RNAP concentration in presence of Cm, (Fig. S3B).

**Estimation of the change in ppGpp concentration from the analysis of the effect of Cm on GFP expression from the P5 and P1 promoters.** P1, as a ribosomal RNA promoter, is inhibited by ppGpp. Therefore, its transcription rate is determined by both its binding affinity with RNAP and ppGpp concentration. ppGpp together with DksA mainly affect the isomerisation to the open complex rather than the promoter binding step (5, 6). The relative transcription rate of P1 can then be approximately formalized as

$$TR(P1) = F(c_p) \cdot c_f/(c_f + K_1), \qquad [6]$$

where $K_1$ is the dissociation constant and $F(c_p)$ is a function of ppGpp concentration ($c_p$). For simplicity, we assume

$$F(c_p) = b \cdot K_p^n/(c_p^n + K_p^n) \qquad [7]$$

where $K_p$ and $n$ are microscopic dissociation constant and Hill coefficient for ppGpp inhibition, and $b$ is a scaling factor.

From Eqs.3 and 6, the ratio of the GFP production rates for P1 and P5 is expressed as

$$\begin{aligned} Gpr(P1)/Gpr(P5) &= TR(P1)/TR(P5) \\ &= b\frac{K_p^n}{c_p^n + K_p^n}\frac{c_f + K_5}{c_f + K_1}. \end{aligned} \qquad [8]$$

So the effect of ppGpp inhibition can be represented by

$$bK_p^n/(c_p^n + K_p^n) = Gpr(P1)/Gpr(P5)/[(c_f + K_5)/(c_f + K_1)]. \qquad [9]$$

In the absence of Cm, ppGpp concentration can be expressed as a function of growth rate, i.e. $c_p = c_{p0}\exp(-\mu/\mu_p))$, where $c_{p0}$ and $\mu_p$ can be fitted with the data of Bremer and Dennis (7): $c_{p0} = 96.6$ pmol/OD$_{460}$ and $\mu_p = 1.06$ doublings/hour. With this function, ppGpp concentration can be estimated at the different experimental growth rates. Since $c_p$ and the right side of Eq.9 are known, we determine $b$, $K_p$ and $n$ by fitting with Eq.9: $b = 4.1$, $K_p = 42pmol/OD460$ and $n = 3.5$. We can then estimate the ppGpp concentration in presence of Cm (Fig. S4).

**Estimation of the change in transcription and translation rates as a function of Cm.** Using the determined $K_5$ and free RNAP concentration into Eq.3, we can estimate the relative transcription rates (promoter activites) for P5. We use the ratio between GFP production rate and relative transcription rate (i.e. $Gpr/TR$) to indicate the relative translation rate. The relative translation rate is the same for all the three promoters according to the assumption, i.e. $Gpr(P5)/TR(P5) = Gpr(PLtet)/TR(PLtet) = Gpr(P1)/TR(P1)$. Then the common relative translation rate can be determined by $translation\ rate = Gpr(P5)/TR(P5)$. Finally, we estimate the transcription rates for PLtet and P1 by $TR(PLtet) = Gpr(PLtet)/translation\ rate$ and $TR(P1) = Gpr(P1)/translation\ rate$.

Fig. S15 shows that the partitioned Cm effects on GFP expression from three promoters at the transcription and translation levels. The differences between the results obtained with the three promoters constructs can be explained by the differences in RNAP affinity and the regulation by ppGpp of P1 transcription.

**A quantitative model of the probability of a stalled ribosome as a function of gene length as a function of chloramphenicol concentration.** Recently, Hwa and coworkers (8) have proposed a qualitative model to describe the effect of ribosome inhibition on protein synthesis which proposes that the stalling of the ribosomes on the mRNA results in an increased rate of mRNA degradation. A quantitative framework was developed to describe this process. We have started from this framework to develop a model to predict the effect of translation inhibition on the number proteins synthesized from a mRNA as a function of gene length and gene position in an operon.

The assumptions we adopt here are the same as Dai *et al.* (8): the first translating ribosome is coupled to RNA polymerase (9), and the consecutive flow of translating ribosomes along mRNA protects mRNA from degradation by RNase enzymes (10). The presence of a stalled ribosome results in an unprotected gap of the mRNA which is exposed to cleavage by RNaseE endonuclease (10–12). Inhibition of ribosome activity by Cm increases the stalling probability. The ribosomes behind the stalled one are released from the mRNA by the tmRNA/ArfA/YaeJ rescue pathways (13), and the newly exposed mRNA is further degraded by a 3'-exonuclease (12). The leading ribosomes that are not stalled complete translation, and the newly exposed mRNA is degraded (12, 14, 15).

Starting from the above sequence of events, we developed a quantitative model to try to explain the observed effects of Cm on protein expression as a function of gene length. Assuming that stalling is a Poisson process with rate $\beta$, the probability that one translating ribosome is stalled before time $t$ is $f_{stall}(t) = 1 - \exp(-\beta t)$ under the initial condition $f_{stall}(0) = 0$. Notice that if the stall of a ribosome is only caused by Cm hit, then the stall probability of a ribosome ($f_{stall}$) is equal to the probability of a ribosome hit by Cm ($P_{hit}$) proposed by Dai et al. (8). The probability that a ribosome is not stalled on a mRNA with length $L$ (i.e. successfully completing translation) is then

$$1 - f_{stall}(L/v) = \exp(-\beta L/v), \quad [10]$$

where $v$ is translation elongation rate. The mRNA here should have a minimal nonzero length ($L_{\min}$), i.e. $L \geq L_{\min}$ since the ORF length close to zero is meaningless. Moreover, the stalling events can be also induced by the intrinsic nature of the mRNA sequence (e.g. the anti-Shine–Dalgarno sequence and pseudoknots) (reviewed in (13)), so the rate $\beta$ has a minimal value in absence of Cm, while the presence of Cm increases it. Thus we represent $\beta$ as a linear function of chloramphenicol concentration ([Cm]) by

$$\beta = \beta_0 + k[Cm] \quad [11]$$

where $\beta_0$ indicates the stalling rate in the absence of Cm. We estimate $\beta_0 = 0.007s^{-1}$ by fitting the data of Tsung et al. (16) (Fig. S5). $L \geq L_{\min}$ and $\beta_0 > 0$ guarantee that $\exp(-\beta L/v) < 1$.

We consider that each translating ribosome has the same stalling probability. Moreover, We assume the stalling of a ribosome causes mRNA degradation with a probability of 100% by the mechanisms described above since the stalling time of a ribosome hit by Cm is usually long because of the strong binding affinity between Cm and ribosome (17). Thus the probability that $n$ proteins are synthesized, in total, from one mRNA with length $L$ (i.e. that the $n+1$th translating ribosome is stalled and induces mRNA degradation and translation abortion of the ribosomes following it) is:

$$p(n) = (1 - f_{stall}(L/v))^n f_{stall}(L/v) = \exp(-n\beta L/v)(1 - \exp(-\beta L/v)) \quad [12]$$

The average number of proteins produced from one mRNA is:

$$\langle n \rangle = \sum_{n=0}^{\infty} n \exp(-n\beta L/v)(1 - \exp(-\beta L/v)) = \frac{1}{\exp(\beta L/v) - 1} \quad [13]$$

Notice that, in principle, $\langle n \rangle$ cannot go to infinity because $L \geq L_{\min}$ (see above).

The ratio of protein production from a shorter mRNA (length $L_1$) to the one from a longer mRNA (length $L_2$) is

$$\frac{\langle n_1 \rangle}{\langle n_2 \rangle} = \frac{\exp(\beta L_2/v) - 1}{\exp(\beta L_1/v) - 1} \quad [14]$$

In order to fit this equation with the experimental data, we need to express $\beta$ and $v$ as functions of growth rate ($\lambda$) or Cm concentration ([Cm]). Under translation limitation, the RNA to protein ratio ($r$) is negatively correlated with growth rate ($\lambda$) by

$$r = r_{max} - \lambda/k_n \quad [15]$$

where $r_{max}$ is the maximal RNA to protein ratio and $k_n$ is the "nutritional capacity" reflecting the quality of the growth medium (18). The ribosome elongation rate ($v$) and $r$ have a Michaelis-Menten relation:

$$v = \frac{v_{max} r}{r + K_M} \quad [16]$$

where $v_{max} = 22a.a./s$ and $K_M = 0.11$ (8, 19). Thus $v$ can be expressed a function of $\lambda$.

Furthermore, we need to estimate the relationship between $[Cm]$ and growth rate. From Dai et al. (8), the fraction of active ribosomes can be expressed as

$$f_{active} = \frac{N_{Rb}^{active}}{N_{Rb}} \propto \frac{\lambda}{v \cdot r} \quad [17]$$

and the ratio of ribosomes not bound with chloramphenicol can be represented by

$$\frac{[Rb]_f}{[Rb]_{tot}} = \frac{K_D}{K_D + [Cm]}, \quad [18]$$

where $K_D$ is the dissociation constant which may also reflect the effects of Cm uptake and other processes. We assume

$$\frac{f_{active}}{f_{active}[0]} \approx \frac{[Rb]_f}{[Rb]_{tot}} \quad [19]$$

where $f_{active}[0]$ indicates the active ribosome fraction in absence of Cm, since active ribosome fraction and free ribosome ratio decreases with Cm concentration similarly (8). Based on Eqs.15-19, we obtain a quantitative relation between growth rate and $[Cm]$

$$\lambda = k_n(r_{max} - 1/F([Cm])), \quad [20]$$

where

$$F([Cm]) = -0.5\left(\frac{1}{K_M} - \frac{1}{r_{max}}\right) + \sqrt{\frac{1}{4}\left(\frac{1}{K_M} - \frac{1}{r_{max}}\right)^2 + \frac{g(\lambda_0)}{k_n r_{max} K_M} \frac{K_D}{K_D + [Cm]}},$$

$$g(\lambda_0) = \lambda_0(r_{max} - \lambda_0/k_n + K_M)/(r_{max} - \lambda_0/k_n)^2,$$

and $\lambda_0$ is the growth rate in the absence of Cm. The dissociation constant $K_D$ for Cm binding with ribosome measured in different labs varies from 0.5 to 5 $\mu M$ (17). So we take $K_D$ as a free parameter and fix it by fitting the empirical growth rate as a function of Cm concentration with Eq.20.

Greulich et al.(20) also derived a formula for growth rate as a function of Cm concentration based on a mathematical model which integrates the dynamics of the uptake and binding with ribosome of chloramphenicol and two empirical relationships of growth rate and ribosome fraction from Scott et al. (18), i.e.

$$[Cm] = \frac{2IC^*_{50}}{\widetilde{\lambda}^*}\left(-\widetilde{\lambda}^2 + \widetilde{\lambda} + \frac{\widetilde{\lambda}^{*2}}{4\widetilde{\lambda}} - \frac{\widetilde{\lambda}^{*2}}{4}\right) \quad [21]$$

where $\widetilde{\lambda} = \lambda/\lambda_0$, $\widetilde{\lambda}^* = \lambda^*_0/\lambda_0$ and $IC^*_{50}$, $\lambda^*_0$ and $\lambda_0$ are parameters dependent on the growth medium.

Eq 14 gives the expression ratio of two genes of different lengths. Putting Eq. 11 into Eq.14, we obtain that the ratio increases with increasing Cm concentration. The equations 14-11 and 20 give a good fit to the experimentally determined expression ratio of GFP to $\beta$-galactosidase as a function of growth rate (see Fig. 3 in the main text and Fig. S6) within reasonable parameter values: $L_{lacZ}(L_2) = 1024\ a.a.$, $L_{gfp}(L_1) = 238\ a.a.$; $v_{max} = 22\ a.a./s$, $K_M = 0.11$ (8, 19); $r_{max}$=0.668, $\lambda_0 = 1h^{-1}$ (18); $\beta_0 = 0.007s^{-1}$ (fitted with the data of Tsung et al. (16)); $k = 0.0088(\mu M \cdot min)^{-1}$, $K_D = 1.5\mu M$, scale factor= 0.216 (fitted with the data of GFP vs lacZ). Replacing Eq. 20 with Eq. 21, the model can also give a good fit (Fig. S7) with two extra parameters (instead of $K_D$): $IC^*_{50} = 4.5\ \mu M$ and $\lambda^*_0 = 1.28\ h^{-1}$ (20).

**A decrease in translation processivity can result in decreased expression of late operon genes.** Transcription and translation are coupled via a physical interaction between RNA polymerase and the first ribosome translating the mRNA, which can be mediated by NusG or RfaH (21–24). Therefore, an additional factor that could decrease the expression of longer genes, and of late genes within an operon, in the presence of translation inhibitors is the loss of this RNAP-ribosome interaction exposing the mRNA for degradation and/or decreasing RNAP speed and processivity (9).

Previous work has shown that inhibiting ribosome activity with a higher concentration of antibiotics than was used here can result in decoupling of transcription and translation (9). To test whether these sublethal concentrations of Cm could have a similar effect on the RNAP-ribosome interaction, constructs were made where two genes of equal length coding for a red and a green fluorescent protein are placed one after the other within the same operon (Figure ??A). In this case there is an equal probability that a ribosome will stall during translation of either gene. However, if the first ribosome translating the upstream gene is inhibited, the one in contact with RNAP, it will also decrease the probability that the downstream gene will be transcribed. Indeed, in these constructs we observe a decrease in the GFP to RFP ratio with increasing Cm. This could be either due to a decrease in transcription processivity from the loss of the RNAP-ribosome interaction or to an increase in the probability that the operon mRNA is degraded before RNAP finishes GFP transcription due to early termination of RFP translation, or a combination of both.

The formation of a terminator hairpin in the transcribed mRNA can be used to detect the presence of a ribosome-RNAP interaction (25). If a ribosome is bound on the mRNA as it is being extruded from RNAP, the terminator hairpin structure cannot fold. If inhibition of ribosome activity decouples translation from the ongoing transcription, then the RNA will be allowed to fold and transcription will stop. Transcription terminators of different strength have been inserted between the two genes (25). The efficiency of these terminators is determined by the distance between the stop codon of the upstream gene and the hairpin loop forming sequence. Termination will be less efficient when this distance is shorter. If transcription and translation are decoupled, the RNAP to ribosome distance will increase and we expect that the efficiency of the weaker terminators will increase with increasing Cm. However, we observe the same fractional decrease in GFP/RFP in all the constructs, independently of the distance from the stop codon, or of the presence of the terminator (Fig. S8). Therefore, it seems that at these low Cm concentrations, the probability of decoupling of transcription and translation is not significant enough to allow for the folding of the hairpin loop. The stalling of a ribosome however, independently of its interaction with RNAP, can result in an increased probability of mRNA degradation. In this case, the stalling of non-leading ribosomes, which are in greater number than the one interacting with RNAP and thus a more probable target, can decrease the lifetime of the operon's mRNA, decreasing the probability that translation will be completed at the *gfp* gene. These results can be reproduced by a model using the parameters values for the probability of translation termination obtained from the comparison in GFP vs LacZ translation (Fig. 3) and the terminator strengths measured in the previous study (25) (Fig. ??B).

**The quantitative model of the probability of a stalled ribosome leading to mRNA degradation can also explain the effect of chloramphenicol on gene expression as a function of gene position in an operon.** The above model of the probability of a stalled ribosome causing a decrease in further gene expression via mRNA degradation can also be used to explain the effects of Cm on gene expression related to the position of the gene in an operon. Because the upstream gene is always transcribed before the downstream gene, at first there will be more ribosomes on the upstream gene than on the downstream one. Moreover, a stalled ribosome on the upstream gene will stop transcription of the downstream gene. Therefore, there is a higher probability that a ribosome stalled on the upstream gene leading to the operon mRNA degradation will affect the expression of the downstream gene than *vice versa*. This only holds for the translation taking place during transcription. Once the operon mRNA is fully transcribed the ribosome density on the two genes will eventually become the same, with a delay that depends on the transcription rate and the translation initiation frequency. At this point there is an equal probability of hitting a RFP or a GFP translating ribosome. The stalling of a ribosome on the downstream gene will also result in the degradation of the operon mRNA, but it will decrease equally the translation of both genes and thus not affect the ratio between the two. The measurement of the ratio of the two proteins leads to the estimation of the probability that a ribosome in the upstream gene is hit before this equilibrium is reached. Here we consider the RFP-GFP system, shown in Fig. **??** in the main text, where the RFP gene is upstream of the GFP gene. The observed decrease in GFP/RFP with increasing Cm shows that ribosome stalling can take place before this equilibrium is reached and that its probability will increase as Cm is increased, as expected.

Let us set L equal to the length of the RFP gene and 2L the length of the whole operon. The probability that a ribosome is not stalled on a mRNA with length $L$ (i.e. successfully completing translation) is $1 - f_{stall}(L/v) = \exp(-\beta L/v)$, where $v$ is translation elongation rate and $\beta = \beta_0 + k[Cm]$ (see above). Notice that $v$ increases with Cm concentration (8) (see Eqs. 15-16, and 20). At a given Cm concentration, $\beta$ and $v$ are fixed, and then the probability function $1 - f_{stall}(L/v)$ is fixed as well.

For simplicity, as done for the single gene case, we ignore the stochasticity in translation initiation and elongation, i.e. we assume the ribosomes bind to RBS and move along mRNA with a constant rate. Since the ribosome binding sites for RFP and GFP in our RFP-GFP operon system are both close to the consensus, we can assume that RFP and GFP have the same translation initiation rate (denoted by $\alpha$) and the average number of translating ribosome on a mature mRNA is more than 1, i.e. $\alpha L/v >> 1$. We still adopt the mRNA degradation mechanism described by Dai et al. (8), just as was done above for a single gene: a translating ribosome hit by Cm will be stalled more than 10 min on average (17); a stalled ribosome will leave a gap in the mRNA, between it and the ribosome translating in front of it, to which RNase can bind to and induce mRNA degradation by the direct entry pathway (10, 11); the ribosomes following the stalled one will be stalled as well and finally all the stalled ribosomes will be released (13); the mRNA region upstream the stall site will be degraded instantaneously from the 3'- to the 5'-terminal by the 3' exoribonucleases (12); the leading unaffected ribosomes downstream of the stall site will finish the translation and the mRNA region behind them will be degraded in a 5'-monophosphorylated end dependent way (12, 14, 15).

We consider the ribosomes translating RFP and GFP as two "queues" and number each ribosome on the queue sequentially. Each stalled ribosome can lead to mRNA degradation. We denote the queue number of the stalled ribosomes as $n_1 + 1$ and $n_2 + 1$, for RFP and GFP respectively. The number of ribosomes that complete the translation of RFP and GFP, i.e. the number of mature RFP and GFP, are thus no more than $n_1$ and $n_2$, respectively. To determine the numbers of mature RFP and GFP the effect of the degradation from one gene coding region on the translation of another gene also needs to be considered.

We first consider the effect of mRNA degradation from a ribosome stalled on the RFP ORF on GFP translation. By the time $n_1$ ribosomes translating RFP have arrived at the end of the RFP ORF, $n_1$ ribosomes have also initiated the translation of GFP (on average). The degradation of the RFP coding region leads to the degradation of the GFP coding region because the free 5'-monophosphorylated end of the mRNA enhances RNaseE activity (14, 15). Thus, mRNA degradation from the RFP coding region limits the number of mature GFP to no greater than $n_1$. Therefore, the number of mature GFP equals $\min(n_1, n_2) \triangleq N_2$. Considering all the possible combinations of $n_1$ and $n_2$, we can derive the average number of mature GFP from one transcript by

$$\begin{aligned}\langle N_2 \rangle &= \sum_{n_1=0}^{\infty} \sum_{n_2=0}^{\infty} p(n_1) p(n_2) \min(n_1, n_2) \\ &= \frac{x^2}{1 - x^2}. \end{aligned} \quad [22]$$

where $p(n) = \exp(-n\beta L/v)(1 - \exp(-\beta L/v))$ (see Eq. 12) and $x = 1 - f_{stall}(L/v) = \exp(-\beta L/v)$ (see Eq. 10). From Eq. 22, the average number of mature GFP from the operon is smaller than that in the case of GFP transcribed in the absence of the RFP coding region and Cm increases this difference. The degradation of mRNA from the upstream RFP region lowers the translation probability of the downstream GFP and the effect is higher with a higher Cm concentration.

Next we consider the effect of the degradation from the GFP ORF on RFP translation. When the $n_2 + 1st$ ribosome translating GFP is stalled, the number of ribosomes that have entered the GFP ORF is $n_2 + k$, where $k$ is an integer from 1 to $m$ ( $\triangleq \alpha L/v$, where $\alpha$ is translation initiation rate, $L$ is gene length, $v$ is translation elongation rate). As mentioned above, $m$ denotes the maximal number of translating ribosomes on the GFP ORF (with constant rates of translation initiation and elongation). The probability of $k$ equal to each integer between 1 and m is the same. Since strong ribosome binding sites are used in our RFP-GFP operon system, we should have $m \gg 1$. The number of ribosomes translating RFP entirely should be the same as the number of ribosomes that have started to translate GFP, i.e. $n_2 + k$. Considering the degradation starting from both the GFP and RFP ORFs, the number of mature RFP should be $\min(n_1, n_2 + k)$ with a fixed $k$. Integrating all the possible values for $k$, the average number of fully synthesized RFP, is $\sum_k (\min(n_1, n_2 + k))/m \triangleq N_1$ (given $n_1$ and $n_2$). With all the combinations of $n_1$ and $n_2$, finally, we can derive the average number of mature RFP from one transcript of the operon by

$$\begin{aligned}\langle N_1\rangle &= \sum_{n_1=0}^{\infty}\sum_{n_2=0}^{\infty} p(n_1)p(n_2)\sum_{k=1}^{m}(\min(n1, n2+k))/m \\ &= \frac{x^2}{1-x^2}\left[1+\frac{1}{x}-\frac{1}{m}\frac{1-x^m}{x(1-x)}\right] \\ &\approx \frac{x}{1-x}, \end{aligned} \qquad [23]$$

where $m \gg 1$ is used in the last derivation. From Eq. 23, the average number of RFP from the operon is approximately the same as in the case of RFP transcribed in the absence of the GFP coding region, suggesting the degradation from the downstream GFP has a small effect on the probability of translation of the upstream RFP.

Based on Eqs. 22 and 23, the ratio of the average numbers of mature GFP and RFP can be obtained as

$$\frac{\langle N_2\rangle}{\langle N_1\rangle} \approx \frac{x}{1+x} = \frac{1}{\exp(\beta L/v)+1} \qquad [24]$$

If there is a terminator between the RFP and GFP genes, we assume its efficiency is $y$. RFP is expressed either from the middle-terminated transcript only including the RFP coding region or from the full transcript including both RFP and GFP coding regions, but GFP is only expressed from the latter transcript. So the ratio of the averaged GFP and RFP numbers is

$$\begin{aligned}\frac{\langle N_2\rangle}{\langle N_1\rangle} &\approx \frac{(1-y)\frac{x^2}{1-x^2}}{y\frac{x}{1-x}+(1-y)\frac{x}{1-x}} \\ &\approx (1-y)\frac{1}{\exp(\beta L/v)+1}. \end{aligned} \qquad [25]$$

When $\beta$ becomes larger due to translation limitation, the ratio $\langle N_2\rangle / \langle N_1\rangle$ decreases independently of the presence of a terminator. Fig. **??** shows that the ratios of the RFP and GFP levels obtained from experimental data can be fit with the above model. The corresponding parameters are: $L_{gfp} = L_{rfp} = 238\ a.a.$; $k_n$=1.32, $r_{max}$=0.670, $\lambda_0 = 0.58h^{-1}$ (18); $v_{max} = 22\ a.a./s$, $K_M = 0.11$, $\beta_0 = 0.007s^{-1}$, $K_D = 1.5\mu M$,$k = 0.0088(\mu M \cdot min)^{-1}$ (the same values as those above obtained from the fit of the data of GFP vs lacZ); scaling factor=7.82 (fitted); terminator efficiencies of ter 1-4 are 0.167, 0.235 0.526, and 0.837, respectively, the same as those measured previously (26).

**Generality of the results.** In order to determine whether the results obtained here, with this small set of promoters and reporter proteins, can be applied to the results obtained on larger datasets, we carried out an analysis of the effect of gene length and operon position on gene expression in the presence of increasing concentrations of Cm on the dataset from Hui and coworkers (27). In their study the amounts of more than one thousand proteins was measured as a function of increasing Cm concentration and under different growth limitations. The proteins were divided in three subsets on whether their concentration increased (R sector), remained the same or decreased (P sector) with increasing translation limitation (Fig. S9).

The first step is to determine how many of the genes in the R sector are influenced directly and indirectly by a change in ppGpp concentration. To do this we used the transcriptomics data on ppGpp dependent regulation from Traxler *et al.* (28). The comparison of these two datasets shows that about 50% of proteins in the R sector are expressed from genes negatively regulated by ppGpp (Fig. S10 A). In order to determine whether this was a direct or indirect regulation, we analyzed the promoter sequences. Promoters that are negatively regulated by ppGpp have a GC-rich sequence between positions -8 and -1, called the discriminator region (29). This analysis shows that the promoters from the subset of genes inhibited by ppGpp indeed tend to have a better match to the discriminator sequence than the others (Fig S10 B). Therefore, ppGpp-dependent transcription regulation can explain the change in expression of a subset of R sector genes.

Promoter sequences can also be analyzed to estimate RNAP affinity using the statistical-mechanical selection method of Berg and von Hippel (1) (see details as above). This analysis shows that the R sector proteins tend to have promoter sequences with a higher affinity for RNAP, with a peak in the distribution below 5 kT (blue, red and purple lines in Figure S11), than those of the other sets, having a wider distribution with a peak at about 5 kT (Fig. S11 A). The presence of a transcription factor can compensate for a low promoter affinity, however regulation by a transcription factor (as defined in RegulonDB (2)) does not change the repartition between R sector and non-R sector promoters. If ribosomal genes are taken out of the analysis the peak below 5 kT is lost (Fig. S11B). The estimated relative affinities of the three promoters used here are shown on the same graph. All three are below 5 kT, among the higher affinity promoters.

We next looked at the protein length distribution in the different protein subsets. The genes in the R sector inhibited by ppGpp have a distribution with a peak below 600 nucleotides (Fig. S12 A, green line). These tend to be for the most part ribosomal proteins or proteins that regulate ribosome activity (Fig S12 C). The length of the *gfpmut2* and *lacZ* genes, as well as of the $\beta$ and $\beta$' subunits of RNAP are shown for reference. The magnitude of the change in the protein concentration as a function of Cm in the Hui *et al.* dataset was measured by the slope of the data in the plot of the change in protein fraction vs growth rate (Fig. S9). The plot of the distribution of the slope values within different windows of gene length shows a more negative slope for the shorter genes. Once the ribosomal genes are taken out of the analysis, the median slope increases, particularly for the genes around 300 bp long (Fig. S12 B,D). The distribution of the slope value as a function of gene length retains the same overall shape independently of whether the promoters are regulated by ppGpp, regulated by transcription factors or constitutive (Fig. S13). In summary, ribosomal proteins tend to be shorter than average, have higher affinity promoters and be negatively regulated by ppGpp, all factors that can lead to increased expression upon translation limitation in different growth media.

Finally, we carried out an analysis of the effect of a gene's position within an operon (Fig. S14). Despite sharing the same promoter region, genes in the same operon do not always belong to the same sector, as measured by the value of the slope in the protein fraction vs growth rate plot (Fig. S14 A, B). Within a given operon, the occurrence of early genes in the R sector and a downstream gene in the P sector is more frequent than the reverse case (binomial one-sided P-value: $3.2\times10^{-5}$) (Fig. S14 C-E). Consistent with the results obtained here with the RFP-GFP construct (Figure **??**).

**The role of the discriminator region in determining changes of gene expression under translation limiting conditions.** We asked how many of the proteins that have been identified as being part of the R-sector may be directly regulated by (p)ppGpp. In a previous study Hui *et al.* used mass spectrometry to measure the change in concentration of about 1000 different proteins in *E. coli* in the presence of different concentrations of Cm (27). They were thus able to identify which proteins change in concentration as R sector proteins and those that belong to the other sectors, because they are induced under different kinds of limitations. In another study Traxler *et al.* used microarrays to measure the changes in mRNA abundance when (p)ppGpp is either absent from the cells or is increased by the addition of serine hydroxamate (28). A Venn diagram shows that there is some overlap between the genes that were negatively regulated by (p)ppGpp and both R-sector and other sector proteins (Figure S10). We next asked how many of the genes in these subsets are regulated by promoters containing a discriminator sequence. We compared the sequence found between -8 and -1 of a gene's promoter with the one of the *rrnB*P1 promoter thus generating a "discriminator score". While all the subsets had a higher score than a set of random sequences, the subset that is at the intersection of the R-sector proteins and the genes showing an inhibitory effect of (p)ppGpp has a higher score. These results indicate that within the R-sector proteins there are both direct and indirect effects of (p)ppGpp on gene expression and that the direct effects are likely due to the presence of a discriminator region at the promoter.

**Discriminator score.** The discriminator score for each available promoter was calculated as follows. The discriminator region required for ppGpp inhibition is a GC-rich sequence between positions -8 and -1 of the promoter sequence relative to the transcription start site (30, 31). The study of Haugen et al. (32) had shown that the C at -7 is very important for the effect of (p)ppGpp, but that identity of the base at the -3 site is not significant. We therefore assumed a consensus sequence for the discriminator region as

| -8 | -7 | -6 | -5 | -4 | -3 | -2 | -1 |
|---|---|---|---|---|---|---|---|
| c | C | c | c | c | - | c | c . |
| g | | g | g | g | | g | g |

Based on this sequence, we scored each promoter by calculating the similarity of one promoter to this consensus sequence at positions -8 to -1. For *rrnB*P1 the score is 1. If a gene has more than one promoter, we take the highest score as its discriminator score.

**Analysis of data obtained from the literature.** We analyzed previously published data from the literature to test the generality of our results. The slopes of the change in protein levels under translation limitation and the sector partition of the proteome were obtained from Hui et al. (27). The lists of the genes inhibited and uninhibited by ppGpp were obtained from Traxlex *et al.* based on their microarray transcription data of the wild type and ppGpp0 (relA-spoT-) strains compared to cells under isoleucine starvation (28). The information on promoter sequences, gene lengths and gene positions in operons was obtained from RegulonDB (2).

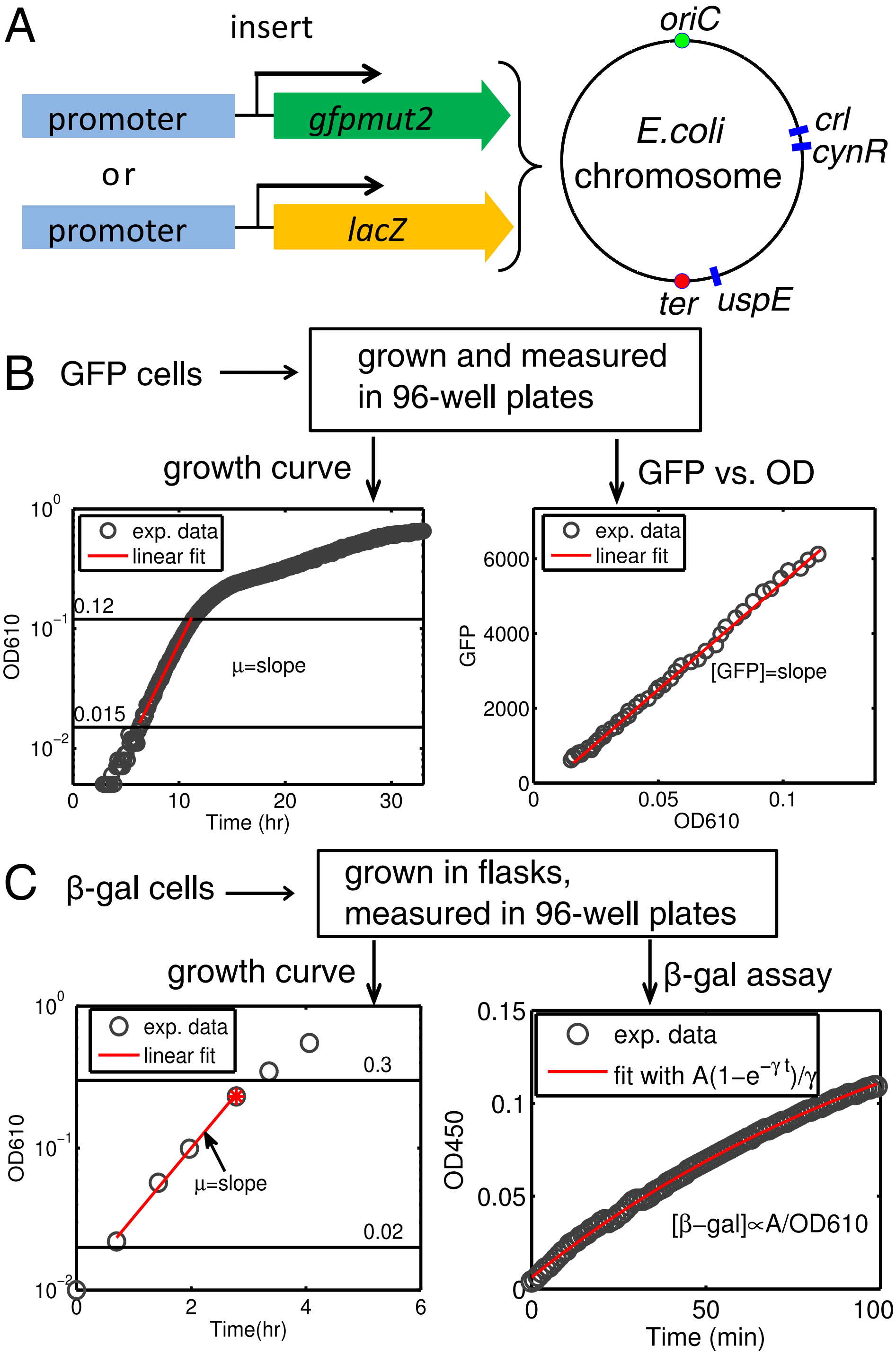


**Fig. S1. Schematic diagram of the experimental designs and data analysis to measure bacterial growth rate and the concentration of the reporter protein.** (A) Chromosomal insertion of the reporter constructs. The reporter gene *gfpmut2* or *lacZ* is fused downstream of a specific promoter (Pltet, P5 or P1) and then inserted in *E.coli* chromosome nearby the gene *crl*, *cynR*, or *uspE*. A kanamycin resistance gene expressed divergently was inserted into the chromosome together with the constructs. (B) The strains carrying GFP were cultivated in a 96-well plate and their optical density (OD) at 610nm and green fluorescence were measured with a plate reader (Tecan, infinite 200Pro) every 7 minutes for 23 hours. The growth rate $\mu$ was derived from the slope of the plot of log(OD) vs. time over the exponential phase, which is defined between a lower and an upper threshold value of OD. GFP concentration ([GFP]) was derived from the slope of the plot of GFP vs. OD over the exponential phase. The comparison of P5-*gfpmut2* with P5-*lacZ* was grown in flasks and its OD and fluorescence were measured with the plate reader every 30-50 minutes. See Li et al. (26) for the RFP-GFP operon system shown in Fig. ??. (C) The strains carrying $\beta$-galactosidase ($\beta$-gal) were grown in flasks and the ODs were measured in 96-well plates with the plate reader. The growth rate was obtained in the same way as for the GFP strains. $\beta$-gal concentration ([$\beta$-gal]) was derived from the 96-well $\beta$-gal assay (see details in Zhang et al. (33)). The red star indicates the time when the culture sample was taken for the beta-galactosidase assay.

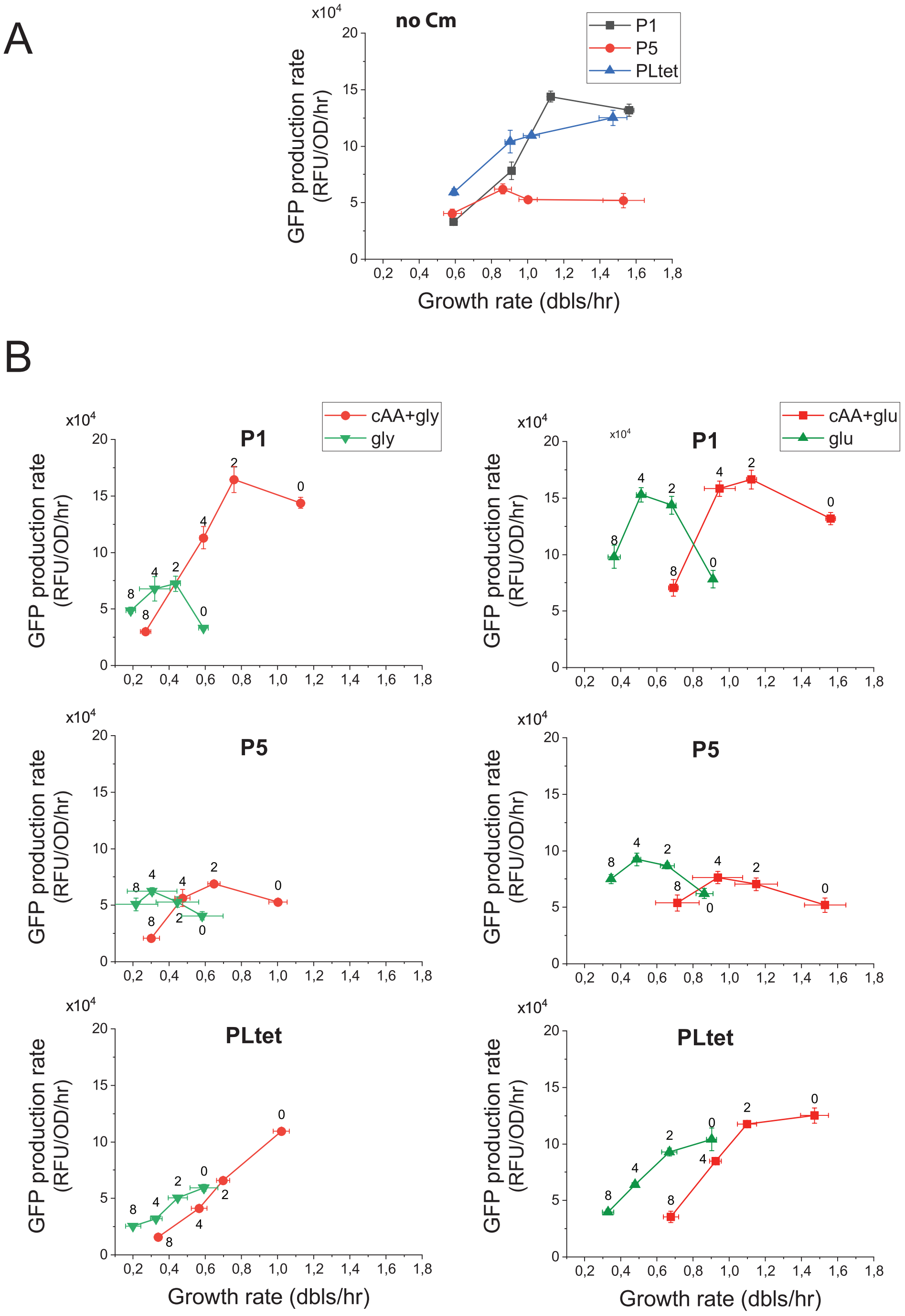


**Fig. S2. Effect of translation limitation on GFP expression from promoters with different affinities for RNAP and different regulation by ppGpp. A.** Change in GFP production rate, as a function of growth rate. Four growth media were used, from the slowest to the fastest: M9-glycerol, M9-glucose, M9-glycerol+casaminoacids, M9-glucose+casaminoacids. P5 and PLtet are both constitutive promoters with different affinities for RNAP while P1 is a shortened version of the rrnBP1 ribosomal RNA promoter with a RNAP affinity similar to P5 but regulated by ppGpp. **B.** Change in GFP production rate as a function of increasing concentration of chloramphenicol in the four growth media. The growth media with casamino acids are in red. The four points correspond to 0, 2, 4 and 8 μM final chloramphenicol concentration as noted next to the data points. The error bars represent the SEM from 3 independent experiments. The error bars smaller than the size of the symbols are not shown. The corresponding changes in GFP concentration are shown in Figure 1 of the manuscript.

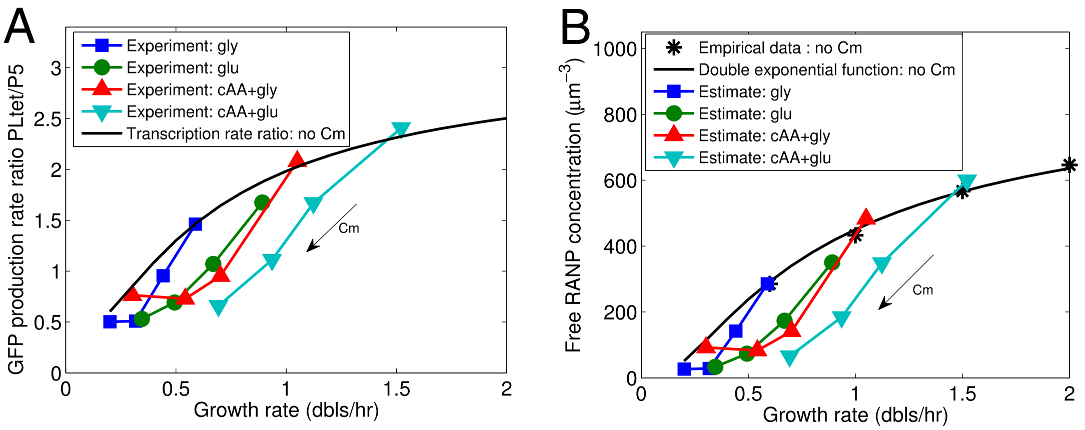


**Fig. S3. Estimated change in free RNAP concentration from the ratio of GFP production rates from the PLtet and P5 promoters.** (A) Measured ratio of GFP production rates from PLtet and P5 as a function of growth rate. The symbols indicate the experimental data. The black line denotes the equation of the ratio of transcription rates from PLtet and P5 without Cm, i.e. $a(c_f + K_5)/(c_f + K_{Ltet})$, where $c_f$ is the free RNAP concentration, $a$, $k_5$ and $K_{Ltet}$ are constants which can be determined by fitting the ratios of GFP production rate to the known values of free RNAP concentrations as a function of growth rate. The free RNAP concentrations as a function of growth rate in the absence of Cm was estimated based on a double exponential function shown by the black line in (B). (B) The black stars correspond to the estimated free RNAP concentration as a function of growth rate obtained from Klumpp and Hwa (4). This data is fit with a double exponential function $c_f = \exp(A\exp(-\mu_r/\mu))$ (black line), where $\mu$ denotes the growth rate, $c_f$ denotes the concentration of free RNAP, $A$ and $\mu_r$ are constants. The free RNAP concentration in the presence of Cm is estimated from the data in (A) with the transcription rate ratio mentioned above. See details of the analysis in the Materials and Methods.

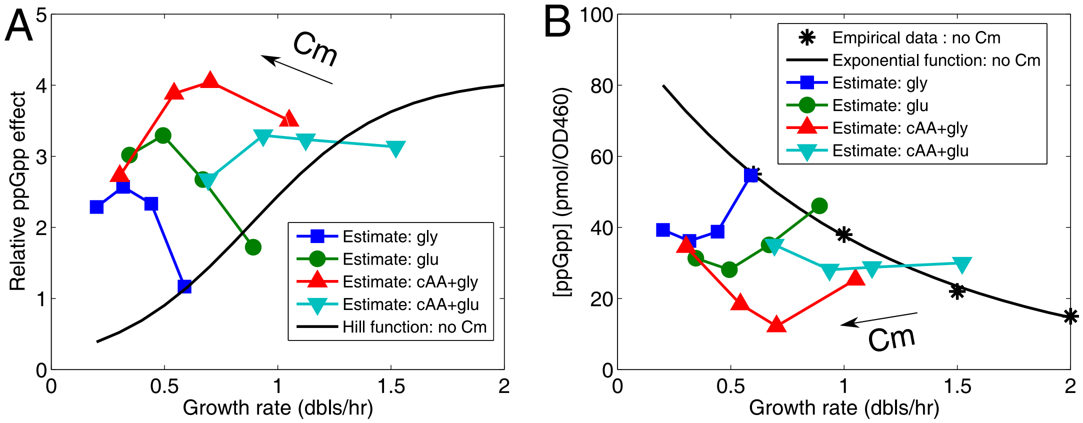


**Fig. S4. Estimated change in the effect of ppGpp on P1 transcription and changes ppGpp concentration as a function of chloramphenicol and growth medium.** (A) The relative ppGpp inhibition effect is estimated as the ratio of GFP production rates from P1 and P5 promoters divided by the ratio of their relative promote activities, i.e. $(K_5 + c_f)/(K_1 + c_f)$, where $c_f$ denotes free RNAP concentration and $K_5$ and $K_1$ are the dissociation constants for P5 and P1 respectively. The black line indicates a Hill function, i.e. $bk_p^n/(k_p^n + c_p^n)$, where $c_p$ is ppGpp concentration, $b$, $k_p$ and $n$ are constants which are fixed by fitting the estimated ppGpp effect without Cm. ppGpp concentration at the growth rates without Cm was estimated based on an exponential function shown by the black line in (B). (B) The black stars correspond to the data on ppGpp concentration as a function of growth rate (7). This data is fit with a function $c_p = c_{p0}\exp(-\mu/\mu_p)$ (black line), where $\mu$ denotes the growth rate, $c_p$ denotes the concentration of ppGpp, $c_{p0}$ and $\mu_p$ are constant. ppGpp concentration in the presence of Cm is estimated from the data in (A) with the Hill function mentioned above. See details of the analysis in the Materials and Methods.

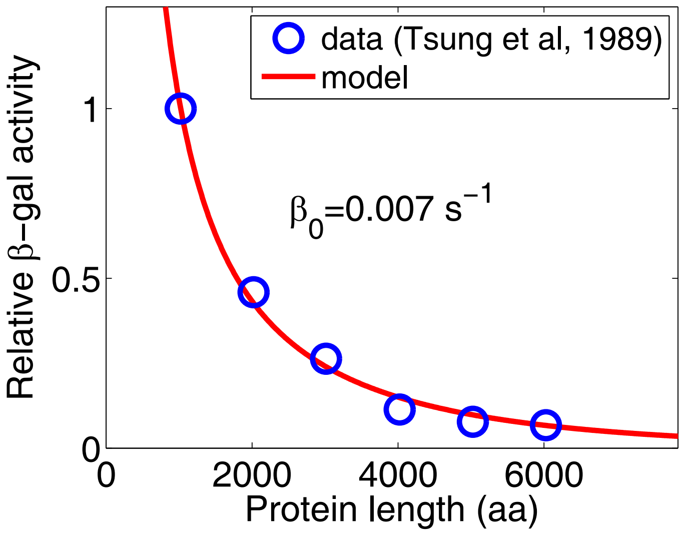


**Fig. S5. Determination of $\beta_0$ by fitting the model with the experimental data of Tsung et al. (16) in absence of Cm.** The proteins of different lengths shown by the circles were expressed from the multimers of the lacZ gene (only the last one confers $\beta$-galactosidase activity). The activities of all the proteins were divided by the activity of wide-type one.

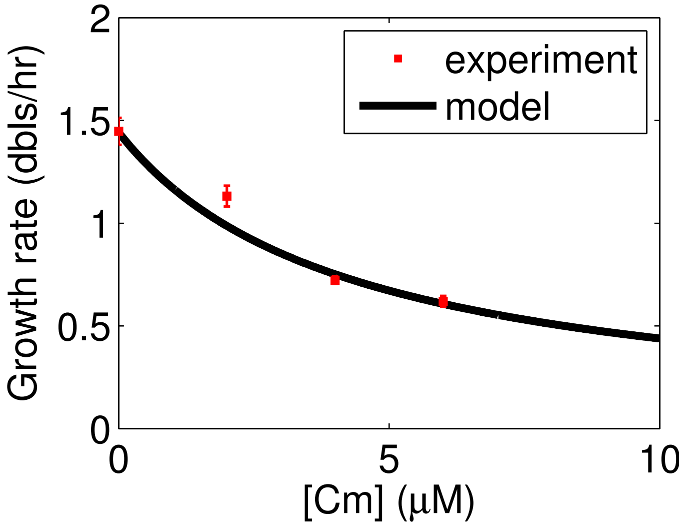


**Fig. S6.** The dependence of growth rate on the Cm concentration can be fit with the formula we derived based on the work of Dai et al. (8).

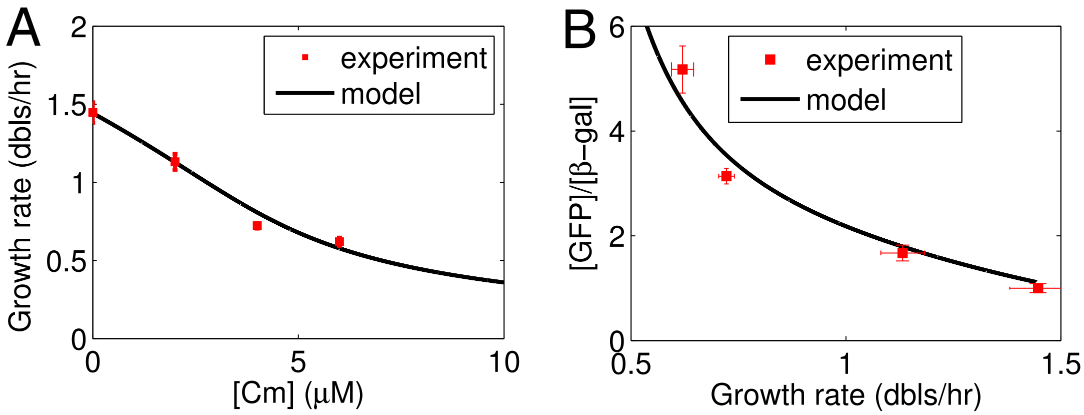


**Fig. S7. The use of formula of Greulich et al. (20) for the relationship between growth rate and chloramphenicol concentration in the model also provides good fits with our experimental data.** (A) The dependence of growth rate on the Cm concentration can be fit with the formula derived by Greulich et al. (20). (B) The concentration ratio between GFP and $\beta$-galactosidase as a function of growth rate can also be fit with the model when the growth rate-Cm formula of Greulich et al. (20) is used.

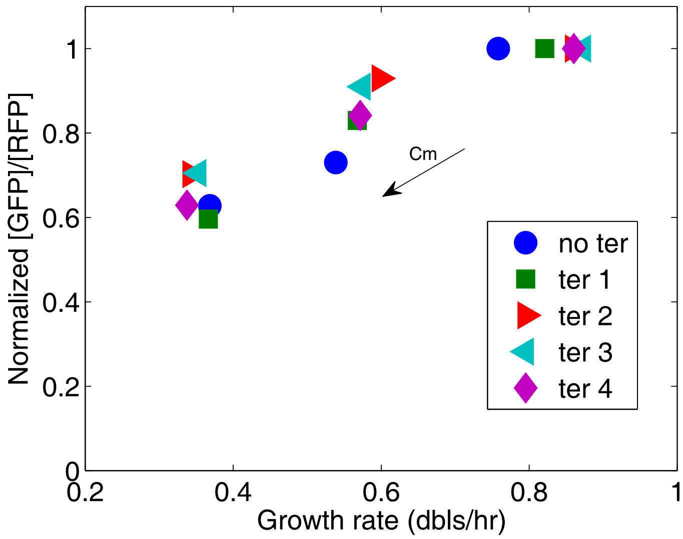


**Fig. S8. Increasing concentration of chloramphenicol result in the same change in GFP/RFP independently of the efficiency of the terminator.** Fractional change in GFP/RFP as a function of growth rate with increasing Cm concentrations (0, 2, 4 $\mu$M), from the data in Fig. ?? (normalized by the point without Cm for each construct).

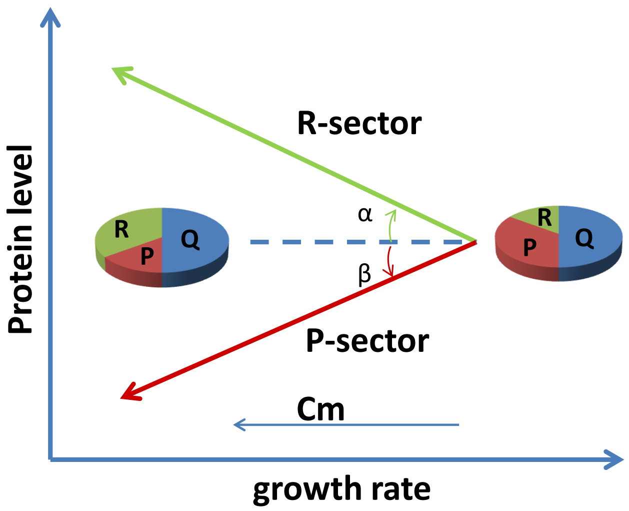


**Fig. S9. Schematic diagram of the divergent change of R-sector and P-sector protein concentrations under translation limitation based on (18).** The difference in the fold change of a given protein as a function of inhibitor can be described by its slope. A negative slope means that the concentration of one protein increases with increasing inhibitor concentration whereas a positive slope means it decreases. The slope for R-sector protein is $-\tan(\alpha) < 0$ and for P-sector protein is $\tan(\beta) > 0$. Q-sector proteins are those whose concentration is not changed by the translation inhibitor. The pies were adapted from Scott et al (18).

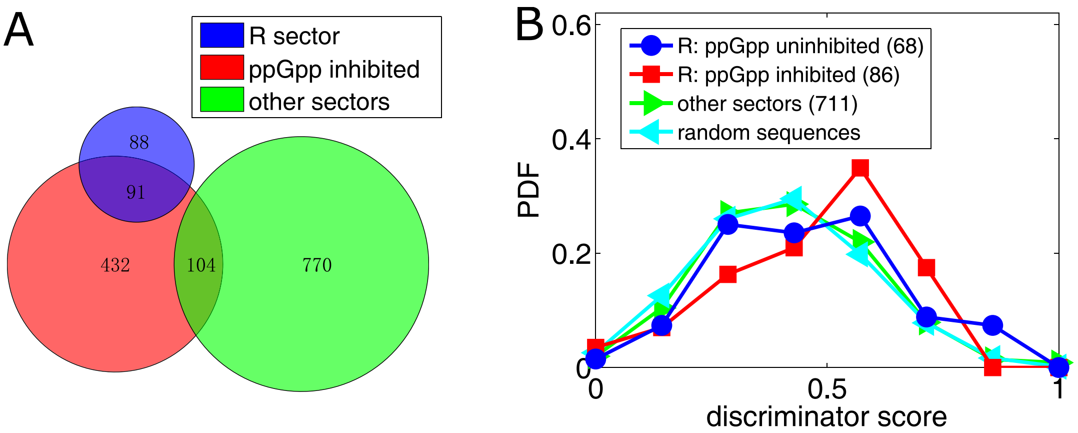


**Fig. S10. A subset of R sector proteins expression is regulated by ppGpp** (A) Venn diagram showing the intersection between the genes whose expression is inhibited (directly or indirectly) by changes in (p)ppGpp concentration (28), R sector proteins and other sector protein identified by mass spectrometry (27). The hypergeometric p-value for over-representation of the intersection between the blue and red sets is $1.5 \times 10^{-32}$, and the p-value for under-representation of the intersection between the red and green sets is 0.012. (B) Distribution of the similarity of the -8 to -1 promoter sequence compared to the consensus sequence of the discriminator region. In producing the distribution of random sequences, the bias in base usage of *E.coli* genome (GC content=43.4%) was considered. The Kolmogorov–Smirnov test for the data set of the R sector inhibited by ppGpp and that of R sector uninhibited by ppGpp plus other sectors gave a two-sided p-value of 0.005.

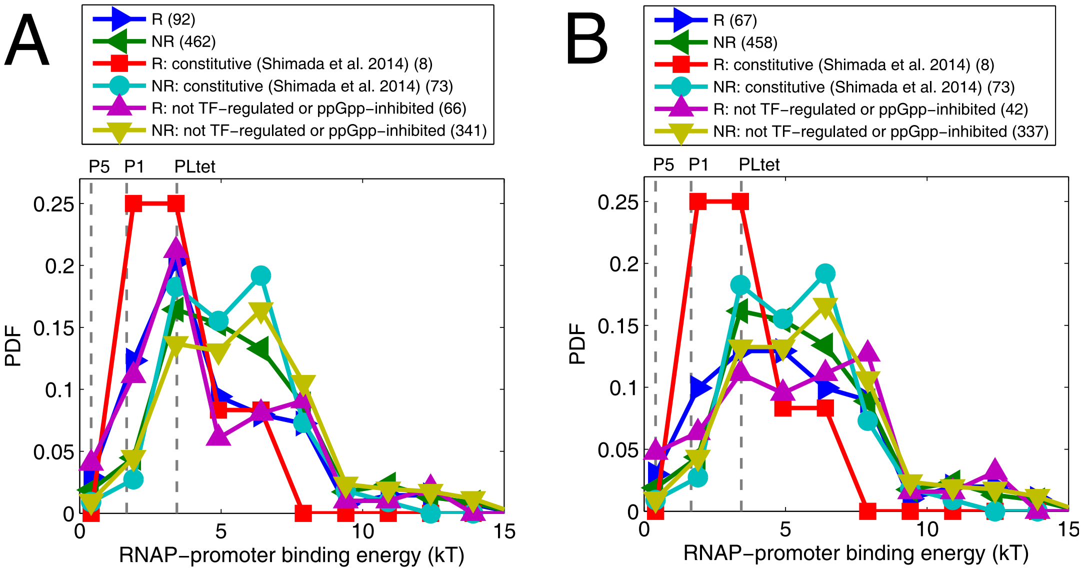


**Fig. S11. Genes whose product belongs to the R sector tend to have a higher affinity for RNAP.** The method of Berg and von Hippel (1) considering spacer length effect was used to estimate promoter affinity for RNAP (see details in the Materials and Methods). The plots compare the RNAP affinities of the promoters in R sector (R) and other sectors (NR) over those all affinity-calculable, wherein Constitutive (as defined by Shimada et al. (34), or wherein not transcription factor(TF)-regulated (based on the regulatory network of all TFs from RegulonDB (2)) or ppGpp-inhibited (based on the transcriptome of Traxler et al. (28)). The number of promoters is shown in parenthesis. (A) Including ribosomal proteins, A gene in R sector tends to have a higher affinity for RNAP than another gene in other sectors over three datasets. (B) Excluding ribosomal proteins. A non-ribosomal gene in R sector tends to have a higher affinity for RNAP than another non-ribosomal gene in other sectors over three datasets. The list of ribosomal proteins are obtained from Ecocyc (35).

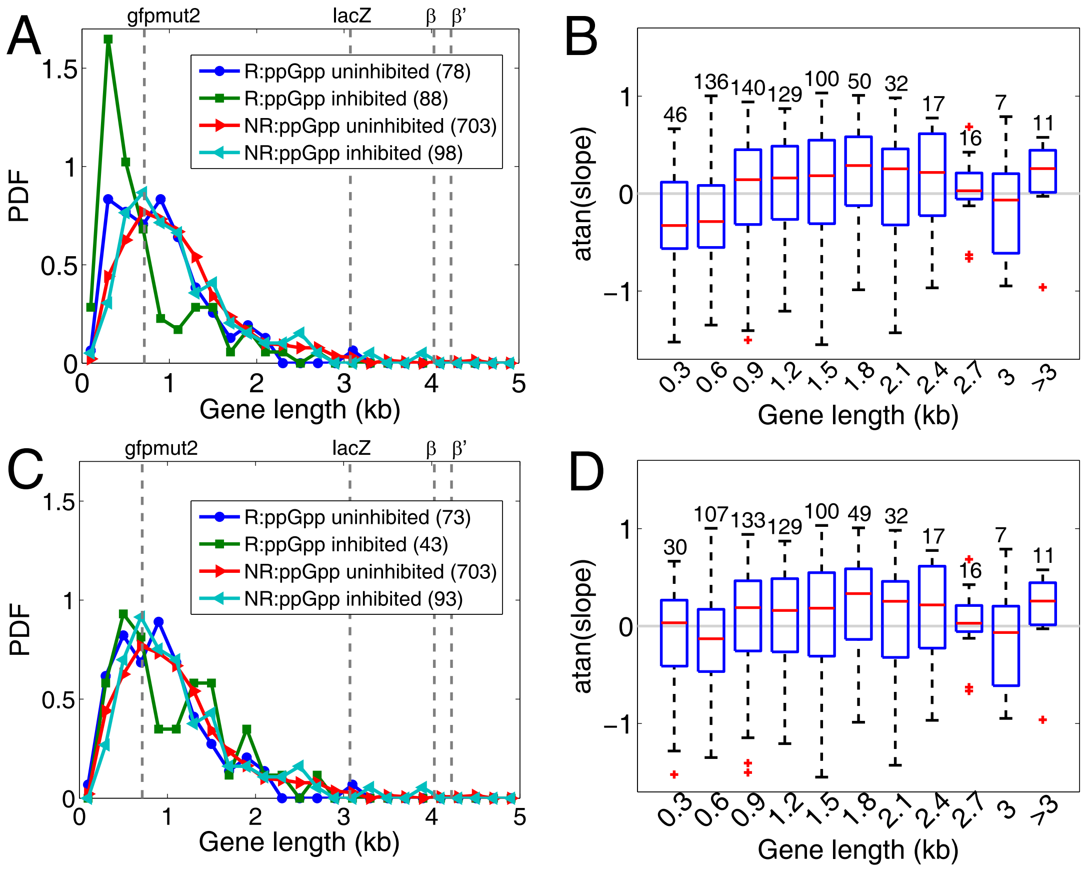


**Fig. S12. Gene length distribution as a function of protein sector and ppGpp regulation.** Most of the short R sector proteins are ribosomal proteins. (A) The sets defined by the intersection of the data of Hui *et al.* (27) and Traxler *et al.* (28) (Fig. S10) were used to calculate the probability density function as a function of gene length. R: R sector proteins, NR: proteins that are not in the R sector. (B) The slope of the change in protein concentration as a function of increasing Cm calculated as in Fig. S9. as a function of gene lengths from the Hui *et al.* dataset (27). The red line is the median, the red cross indicates the degree of significance of the difference with respect to the dataset on the right. C and D show the same analysis without taking into consideration the set of ribosomal proteins (obtained from Ecocyc (35))

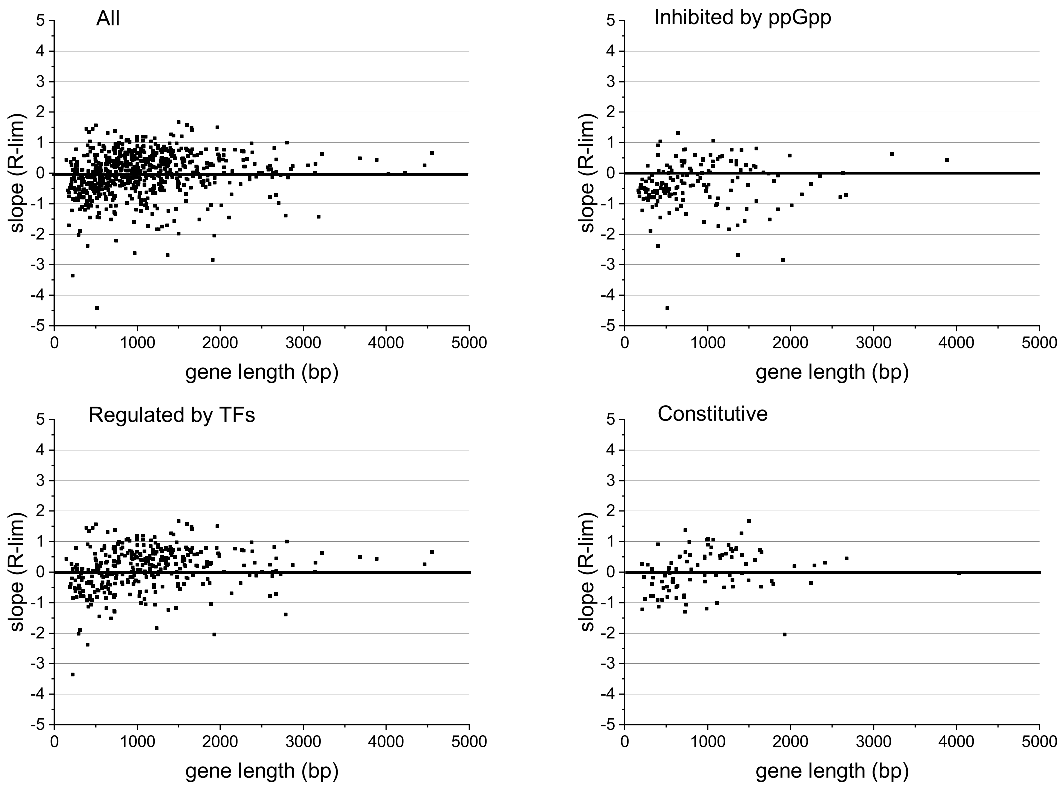


**Fig. S13. The decrease in slope (defined in Fig. S9) with decreasing gene length is independent of whether the gene's promoter is constitutive, regulated by ppGpp or among the set of promoters with known transcription factors (Regulon DB).** The values of the slopes are from Hui et al. (27).

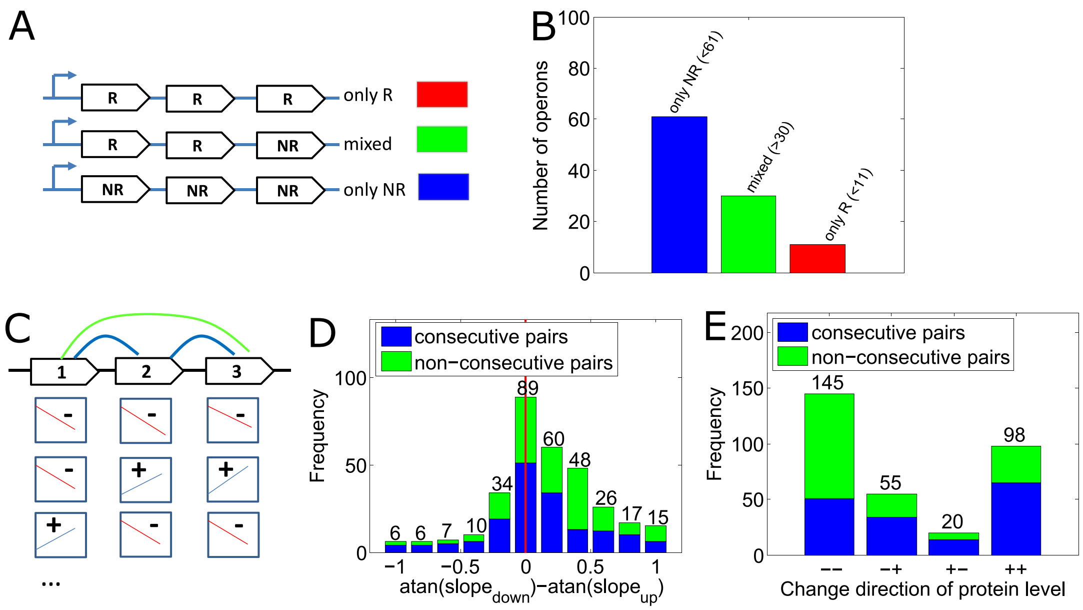


**Fig. S14. A gene's operon position can have an effect on the change in expression upon translation limitation by Cm.** (A-B) Gene operon composition as a function of protein sector. R indicates the R sector, NR indicates the other sectors. (C) Difference in the slope of protein concentration as a function of Cm between two genes of the same operon. The slope of the upstream gene is subtracted from the one of the downstream gene. Note: slope values re not available for all genes in an operon. (D-E) Frequency of change in slope sign between genes in the same operon. The sector partition and slopes used in plots (A,C,D) are from Hui et al. (27).

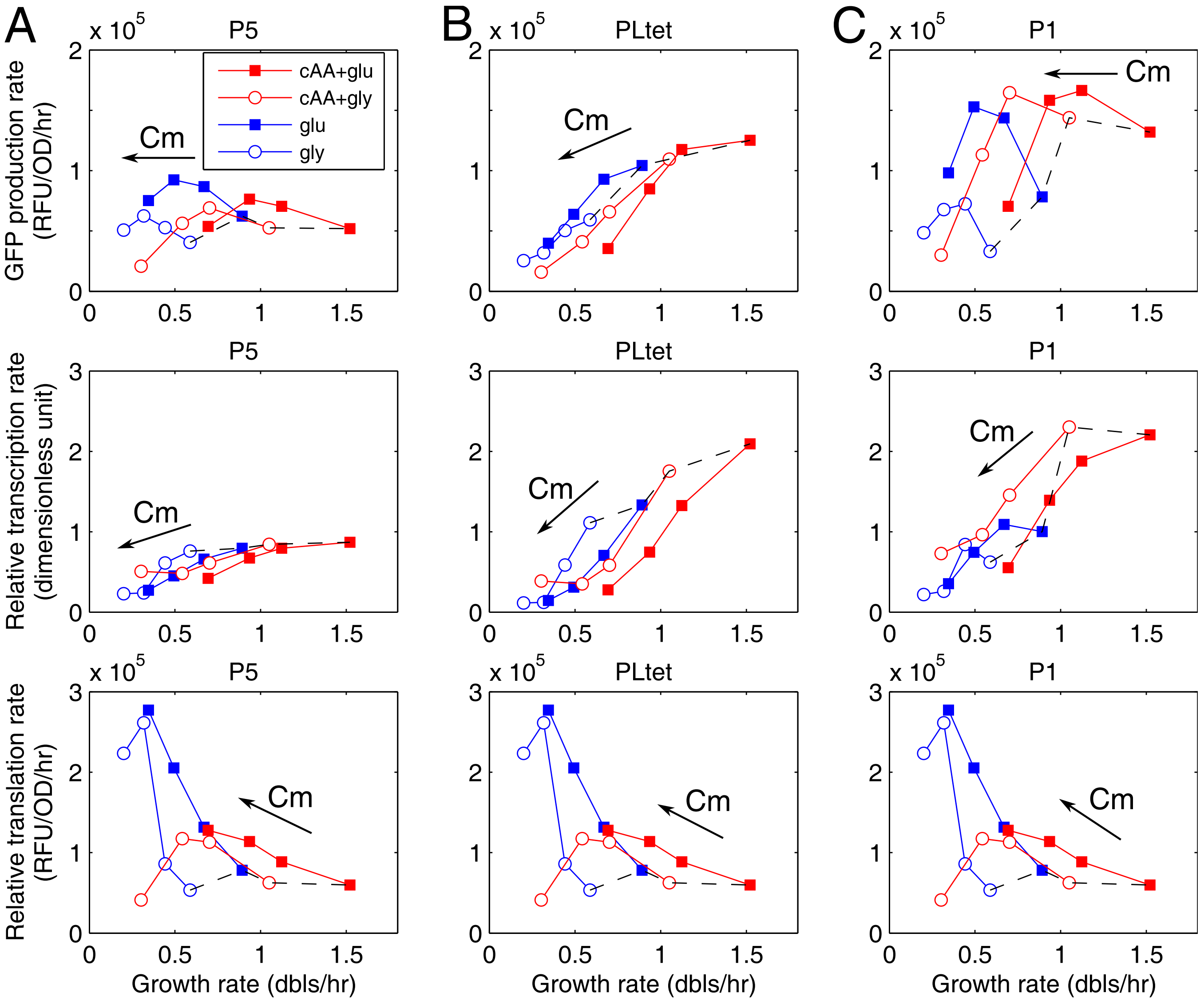


**Fig. S15. Estimation of the change in transcription and translation rate of the *gfpmut2* gene for each promoter construct.** The first row is the same data as in Fig. 1 in the main text. The second row is the (relative) transcription rate (promoter activity) and the third row is the (relative) translation rate. Here we assume that the translation rate for the GFP is the same independently of the promoter. The change in the amount of free RNAP was estimated from the ratio of PLtet to P5 as in Fig. 2 in the main text. This was used to estimate the transcription rate of P5-GFP expression from the dissociation constant $K_5$. The transcription rate was then used to estimate the translation rate that results in the GFP expression rate data in the first row. This translation rate was then used to estimate the transcription rate for PLtet and P1. A similar result is obtained by estimating the transcription rates for PLtet and P1 from their RNAP-promoter dissociation constants.

**Table S1. Sequences of the promoters used in this study. The underlined sites are the -35 and -10 regions. The bold base is the transcription start site. The bases in red in P1 sequence indicate the discriminator region.**

| Promoters | sequences |
|---|---|
| P1 | TTGCGCGGTCAGAAAATTATTTTAAATTTCCTCTTGTCAGGCCGGAATAACTCCCTATAATGCGCCACC**A**CTGACA |
| P5 | ——————ACAACATCTAAGAGAAAAATTATATTGACATCTGCCCTTGAATAAGCTATAATAGTAGT**C**TTAGTTAGAGAAGGAGGGTATAA |
| PLtet | ————————TCCCTATCAGTGATAGAGATTGACATCCCTATCAGTGATAGAGATACTGAGCAC**A**CACAGGAAACA |
